# Modelling white matter in gyral blades as a continuous vector field

**DOI:** 10.1101/2020.07.27.222778

**Authors:** Michiel Cottaar, Matteo Bastiani, Nikhil Boddu, Matthew Glasser, Suzanne Haber, David C. van Essen, Stamatios N. Sotiropoulos, Saad Jbabdi

**Affiliations:** Wellcome Centre for Integrative Neuroimaging (WIN) - Centre for Functional Magnetic Resonance Imaging of the Brain (FMRIB), University of Oxford, UK; Sir Peter Mansfield Imaging Centre, School of Medicine, University of Nottingham, UK; Department of Neuroscience, Washington University Medical School, Saint Louis, MI, 63110, USA; Department of Radiology, Washington University Medical School, Saint Louis, MI, 63110, USA; St. Luke’s Hospital, Saint Louis, MI, 63017, USA; McLean Hospital, Harvard Medical School, Belmont, United States; Department of Pharmacology and Physiology, University of Rochester School of Medicine & Dentistry, Rochester, United States

## Abstract

Many brain imaging studies aim to measure structural connectivity with diffusion tractography. However, biases in tractography data, particularly near the boundary between white matter and cortical grey matter can limit the accuracy of such studies. When seeding from the white matter, streamlines tend to travel parallel to the convoluted cortical surface, largely avoiding sulcal fundi and terminating preferentially on gyral crowns. When seeding from the cortical grey matter, streamlines generally run near the cortical surface until reaching deep white matter. These so-called “gyral biases” limit the accuracy and effective resolution of cortical structural connectivity profiles estimated by tractography algorithms, and they do not reflect the expected distributions of axonal densities seen in invasive tracer studies or stains of myelinated fibres. We propose an algorithm that concurrently models fibre density and orientation using a divergence-free vector field within gyral blades to encourage an anatomically-justified streamline density distribution along the cortical white/grey-matter boundary while maintaining alignment with the diffusion MRI estimated fibre orientations. Using *in vivo* data from the Human Connectome Project, we show that this algorithm reduces tractography biases. We compare the structural connectomes to functional connectomes from resting-state fMRI, showing that our model improves cross-modal agreement. Finally, we find that after parcellation the changes in the structural connectome are very minor with slightly improved interhemispheric connections (i.e, more homotopic connectivity) and slightly worse intrahemispheric connections when compared to tracers.

## 2 Introduction

By tracing continuous paths along the distributions of axonal fibre orientations estimated for each voxel of the brain, diffusion MRI (dMRI) tractography aims to infer the trajectories of white matter fibre bundles. This technique has been used to map the paths of major tracts coursing through white matter or to estimate the connectivity between grey matter regions. This connectivity is often expressed as a “structural connectome”, which is a matrix that contains area to area non-invasive estimates of anatomical connectivity (Sporns, Tononi, and Kötter 2005). Estimating accurate connectivities in the cortex is limited however, by the strong bias of tractography streamlines to avoid sulcal fundi and walls and instead to terminate on gyral crowns and has been termed a “gyral bias” (Van Essen et al. 2014; Reveley et al. 2015; Schilling et al. 2018; Sotiropoulos and Zalesky 2019). This gyral bias limits the accuracy and spatial resolution at which the termination points of white matter bundles can be localised or of grey matter to grey matter connection strength estimation. For example, tractography may localize the termination zone of a streamline to an entire gyrus but not accurately assign it to either sulcal wall, instead leaving it to terminate on the gyral crown. Importantly, we do expect some preference for axons to terminate in the gyral crowns based on the geometry of the gyri (i.e., because of their convex surface curvature, gyral crowns have a greater ratio of overlying grey matter volume to their white matter surface area, sulcal walls have a neutral ratio, and concave sulcal fundi have a smaller overlying grey matter volume to white matter surface area ratio (Van Essen et al. 2014)). However, the gyral bias observed in tractography is much larger than that expected from the geometry of the gyri.

The gyral bias may reflect the strong tendency for fibre bundles to run parallel to the white/grey-matter boundary in the superficial white matter, because even high resolution dMRI fails to adequately capture the sharply curved trajectories of axons ‘peeling off’ to connect with grey matter sulcal walls and fundi seen with histology (Van Essen et al. 2014).

U-fibres connecting neighbouring gyri may be a large contributor to these bundles (Reveley et al. 2015). These parallel fibres lead to the fibre orientations estimated from dMRI to closely align with the nearby white/grey-matter boundary in superficial white matter along sulcal walls and fundi (Figure 1A; Cottaar et al. 2018). Hence most tractography streamlines keep running parallel to the sulcal walls and fundi until they reach the gyral crown, which results in the gyral bias (Figure 1B; Sotiropoulos et al. 2016). While seeding from the cortical volume can reduce this gyral bias by ensuring a uniform distribution of seed points within the cortex, this now creates a bias in the streamline density in white matter, with most streamlines remaining close to the gyral wall (Figure 1C; Smith et al. 2012). Moreover, the gyral bias problem persists for the eventual grey matter terminations of these streamlines as tractography will still overestimate their terminations on gyral crowns versus sulcal fundi. Here we propose a model for gyral white matter that aims to both reduce the overestimation gyral streamline terminations relative to sulcal terminations and the bias of streamlines seeded in the sulcal walls to remain close to the sulcal walls. For our target streamline density distribution when counting on cortical surfaces, we make the first-order assumption that the density of streamline crossing the white/grey-matter boundary in any cortical region should be proportional to the cortical volume divided by the underlying white matter surface area (Van Essen et al. 2014). Thus, our fundamental assumption is that the cortical streamline density per unit cortical volume is uniform; accordingly, we will display our results normalized to unit cortical volume.

**Figure 1.**
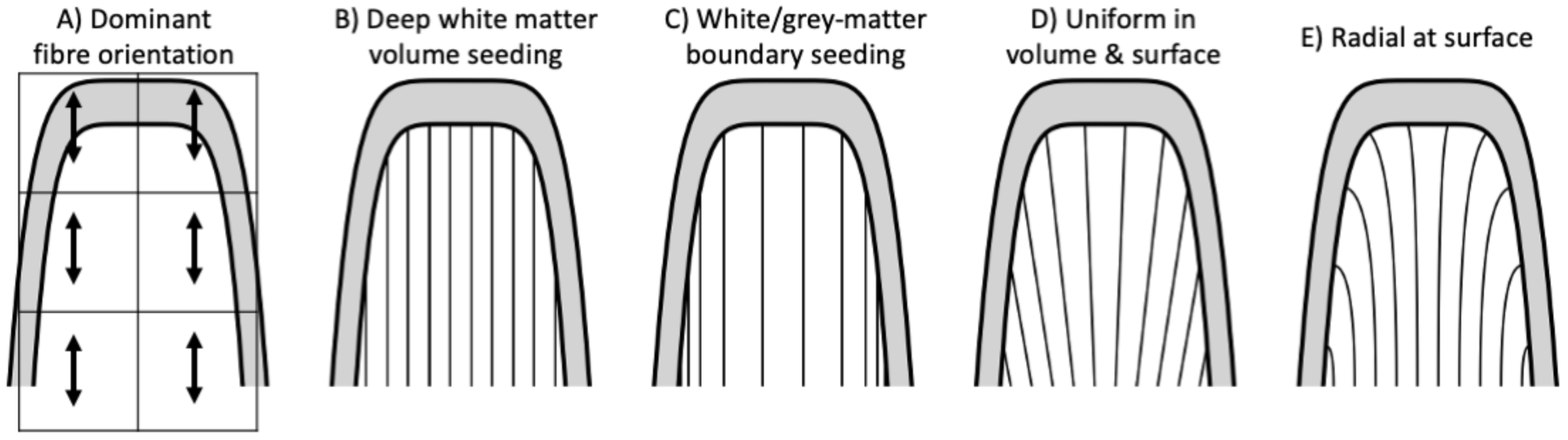
Sketch of possible fibre configurations in white matter of a gyral blade (represented by 10 streamlines). A) Typically, the dominant voxel-wise fibre orientation estimated from diffusion MRI is closely aligned with the gyral wall and points to the gyral crown. This causes two types of gyral biases (B and C): B) It causes local tractography streamlines uniformly entering from deep white matter to preferentially terminate in the gyral crown resulting in a biased density in the cortex. C) Similarly, local tractography streamlines uniformly seeded from the cortex tend to remain close to the gyral walls, resulting in a biased density in the white matter. Note that these streamlines are uniform per associated unit of volume of cortical grey matter rather than uniform across the white/grey-matter boundary. D) By enforcing uniform densities both for the streamlines entering the gyral white matter and in the cortex, more realistic fibre configurations can be obtained, (E) especially if additional constraints such as radiality at the surface are added. Note that the fibre configurations in panels B, C, and E (but not D) are all consistent with the diffusion MRI orientations in panel A.

Our model aims to find a fibre configuration consistent with the diffusion MRI data that has both a uniform density in the white matter within gyral blades as well as a uniform distribution of fibre end-points within the cortical grey matter volume (Figure 1D,E). This requires not only constraining the streamline orientation, but also its density. Hence, we can no longer model a single streamline at a time as in local tractography, but instead need to model the complete set of streamlines at once. To make our initial formulation tractable, we assume that streamlines within gyral blades do not cross or intermix. This means we cannot reconstruct fibres crossing from one side of a gyral bank to the other, which need to be estimated in a different way. Given this assumption, the resultant density constraints create a fibre configuration where the streamlines entering the gyrus at the left will connect to the left gyral wall, while those entering on the right connect to the right gyral wall and those in the centre continue upwards towards the gyral crown (Figure 1D). More realistic 3D fibre configurations can be created by adding additional constraints such as having radial fibre orientations when they reach the cortex (Figure 1E) and alignment with the fibre orientations estimated from diffusion MRI. With this set of geometric and anatomical constraints, the streamlines disperse towards the surface qualitatively similar to that seen in histology (Budde and Annese 2013; Van Essen et al. 2014) and high-resolution diffusion MRI data (Miller et al. 2011; Heidemann et al. 2012; Sotiropoulos et al. 2016).

## 3 Gyral white matter model

### 3.1 Defining gyral white matter

We split the white matter into gyral white matter, which is the white matter contained within the gyral blades, and deep white matter. For the gyral white matter we propose a novel tractography algorithm to describe the white matter configuration not as individual streamlines, but as a continuous vector field. This algorithm is likely to be most accurate in regions where the white matter fibre configuration (i) is constrained by the geometry of the cortical folds (which is typically neglected in local tractography approaches) and (ii) can be accurately described using only single dominant fibre population filling up the available space. While this may be a reasonable description of the white matter over much of the gyral blades, deep white matter is not generally well described in this way. Hence, we implemented a way to apply the novel tractography model to the gyral blades and use standard probabilistic tractography algorithms in the underlying deep white matter.

To define the boundary between the gyral and deep white matter, we introduce a new “gyral thickness” measure for each voxel. This measure is defined as the length of the shortest straight line through the voxel connecting the white/grey-matter boundary on both sides (Figure 2A). This measure is small between the neighbouring sulcal walls and fundi, but large for any white matter below the sulcal fundi. The gyral white matter is any white matter with a gyral thickness less than some threshold. There tends to be a sharp increase in gyral thickness just below the sulcal fundi, so the boundary location is not very sensitive to the exact value chosen for the gyral thickness threshold (Figure 2B). In this study we adopt a threshold of 10 mm.

**Figure 2.**
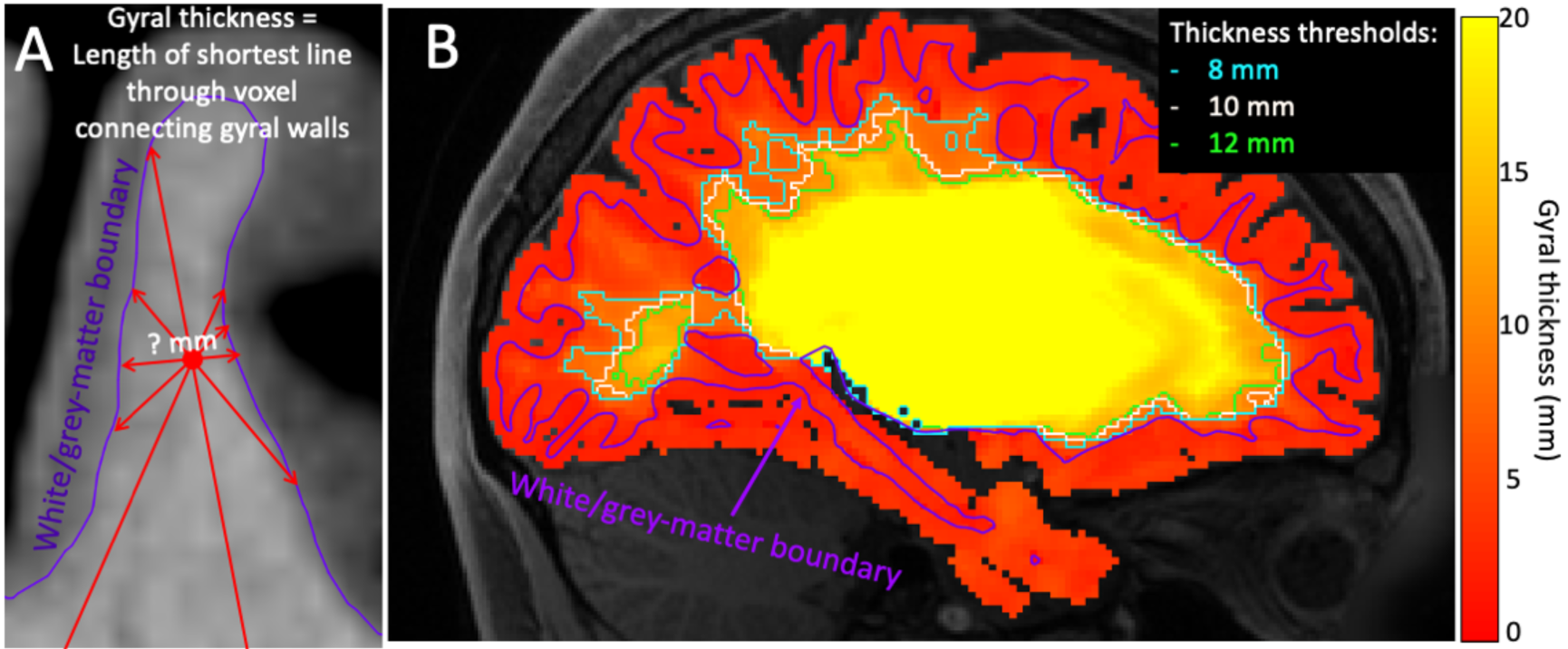
Definition of deep/gyral white matter interface. A) “gyral thickness” is defined as the length of the shortest line through the voxels that hit the white/grey-matter boundary at both ends. B) Thresholding the “Gyral thickness” map separates the white matter in the gyral blades from the deep white matter underneath.

### 3.2 Gyral white model overview

Within the gyral blades we model the fibres as a continuous vector field 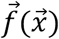. The norm of this vector field 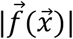 defines the local fibre density at position 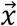, and the orientation of this vector 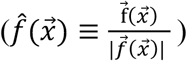 defines the local fibre orientation. Hence, this model allows us to represent, and potentially impose constraints on, both fibre density (e.g., uniform density across the cortex) and fibre orientation (e.g., matching the voxel-wise fibre orientations observed from diffusion MRI).

To produce a realistic fibre configuration, an important constraint is that fibres avoid terminating in white matter. This is strictly enforced by constructing the vector field to be divergence-free:

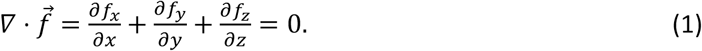

Setting the divergence to zero implies that any decrease in the number of fibres travelling in one direction must be compensated by an increasing number of streamlines in another direction, so that the total number of streamlines traveling along a tract remains constant. This ensures that no streamlines terminate in the white matter.

This single vector field only defines a single fibre orientation and density at every (infinitesimally small) point 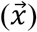 in the brain and hence does not allow for crossing fibres. Crossing fibres could be modelled by describing each white matter tract as a different vector field, which can overlap. However, in this initial formulation we model the superficial white matter as a single, divergence-free vector field, hence ignoring any crossing fibres within this region. Fibre crossings are taken into account for deeper white matter, where we use probabilistic tractography based on a crossing-fibre model (Behrens et al. 2007; Jbabdi et al. 2012), as available in FSL.

Figure 3 shows an overview of our tractography algorithm for the gyral white matter. An initial estimate of the fibre configuration is provided by distributing negative charges at each centre of the triangles in the pial surface mesh and a single, positive charge in deep white matter. The field at any point 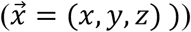 in the white matter is hence given by:

**Figure 3.**
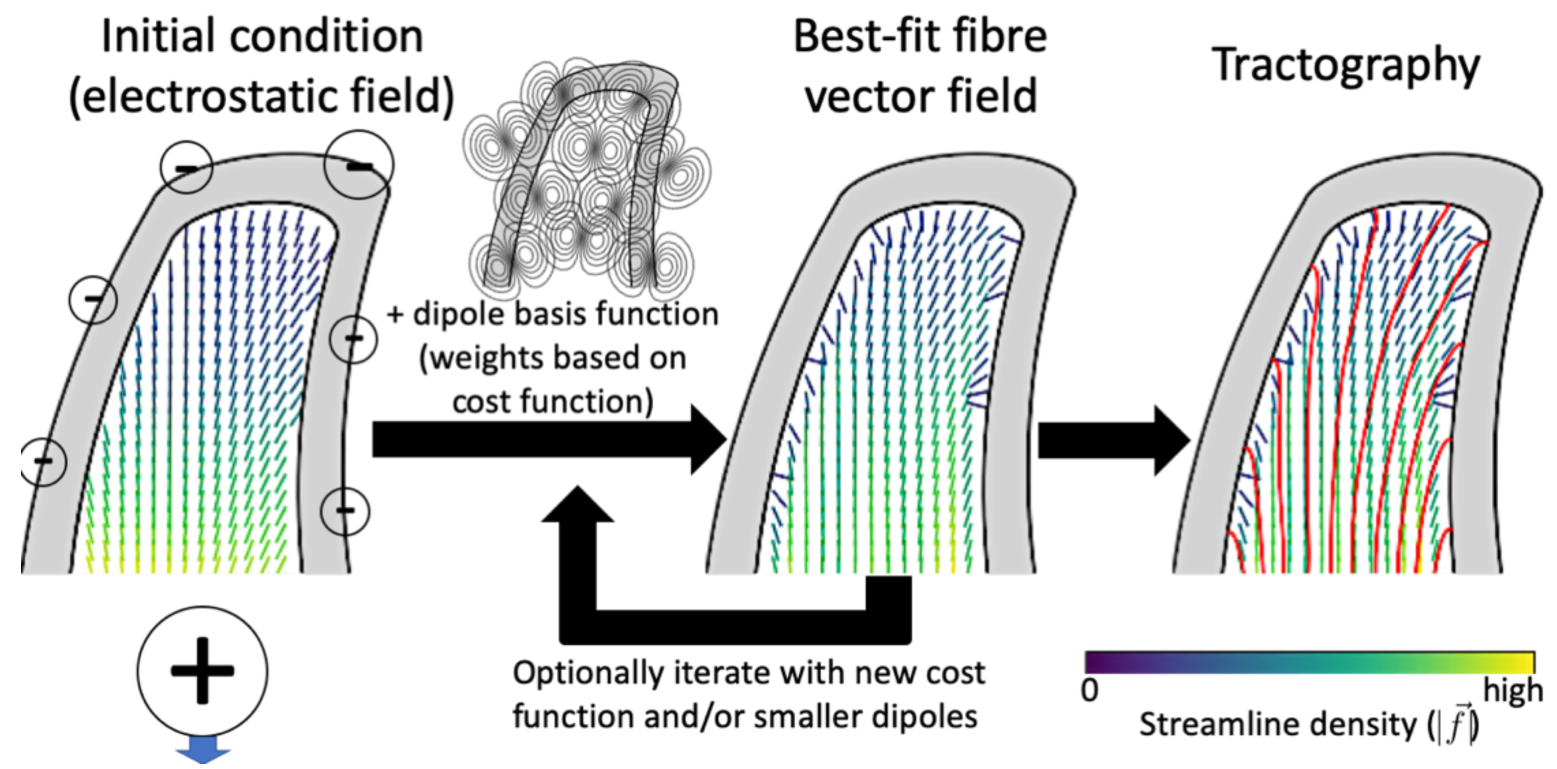
Procedure for modelling the gyral white matter. An initial vector field is estimated from negative electrostatic charges at the pial surface and an equal positive charge in the deep white matter (left). This initial vector field is updated by adding dipole basis functions, where the dipole strengths and orientations are determined by minimizing a cost function, which imposes data fidelity (on fibre orientations) and anatomical constraints (on fibre density and orientation). This step may be iterative with an updated cost function and/or smaller dipoles as basis functions. The resulting vector field configuration can then be used for tractography within the gyral white matter (right). The vector colour encodes the streamline density (colourbar in lower right). The individual steps are explained in sections 3.3-3.4.

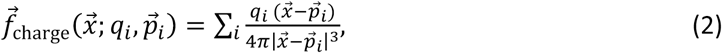

where the *p*_*i*_’s are the positions of the point charges and the *q*_*i*_’s are the charge at point *p*_*i*_. The negative charges at the pial surface are set proportional to the cortical volume represented by that triangle (Winkler et al. 2010). These charges generate streamlines proportional to the cortical volume. The single positive charge in deep white matter is set to the negation of the sum of all negative charges and hence acts as the other termination point for streamlines generated at the pial surface. Note that the resulting vector field is divergence-free, except at the charge locations.

These charges ensure an initial vector field through which streamlines run from deep white matter up to the cortical surface, but the streamlines are not constrained to respect the observed diffusion data or even to remain within the white matter. This vector field is then adjusted while remaining divergence-free by adding a linear combination of dipole-like basis functions (see section 3.3), whose orientation and strength is determined by fitting to a predefined cost function describing constraints on the fibre density and orientation (see section 0). For the example in Figure 3, the cost function encourages both a uniform density distribution along the white/grey-matter boundary and a radial orientation at this surface. The best-fit vector field is used to guide tractography streamlines through the white matter in the gyral blades (see section 3.4).

### 3.3 Dipole basis functions

While streamlines travelling through the vector field generated by the charges defined in section 3.2 will tend to travel from the positive charge in deep white matter to the negative charges along the pial surfaces, they are not constrained to align with the fibre orientations estimated from dMRI or even to traversing through the white matter. By adding divergence-free dipole basis functions to the initial vector field, we can adjust the path of these streamlines (Figure 4) to make them more realistic. In this section we examine these dipole basis functions; in section 0 we investigate the various terms in the cost function used to optimise the dipole orientations and strengths.

**Figure 4.**
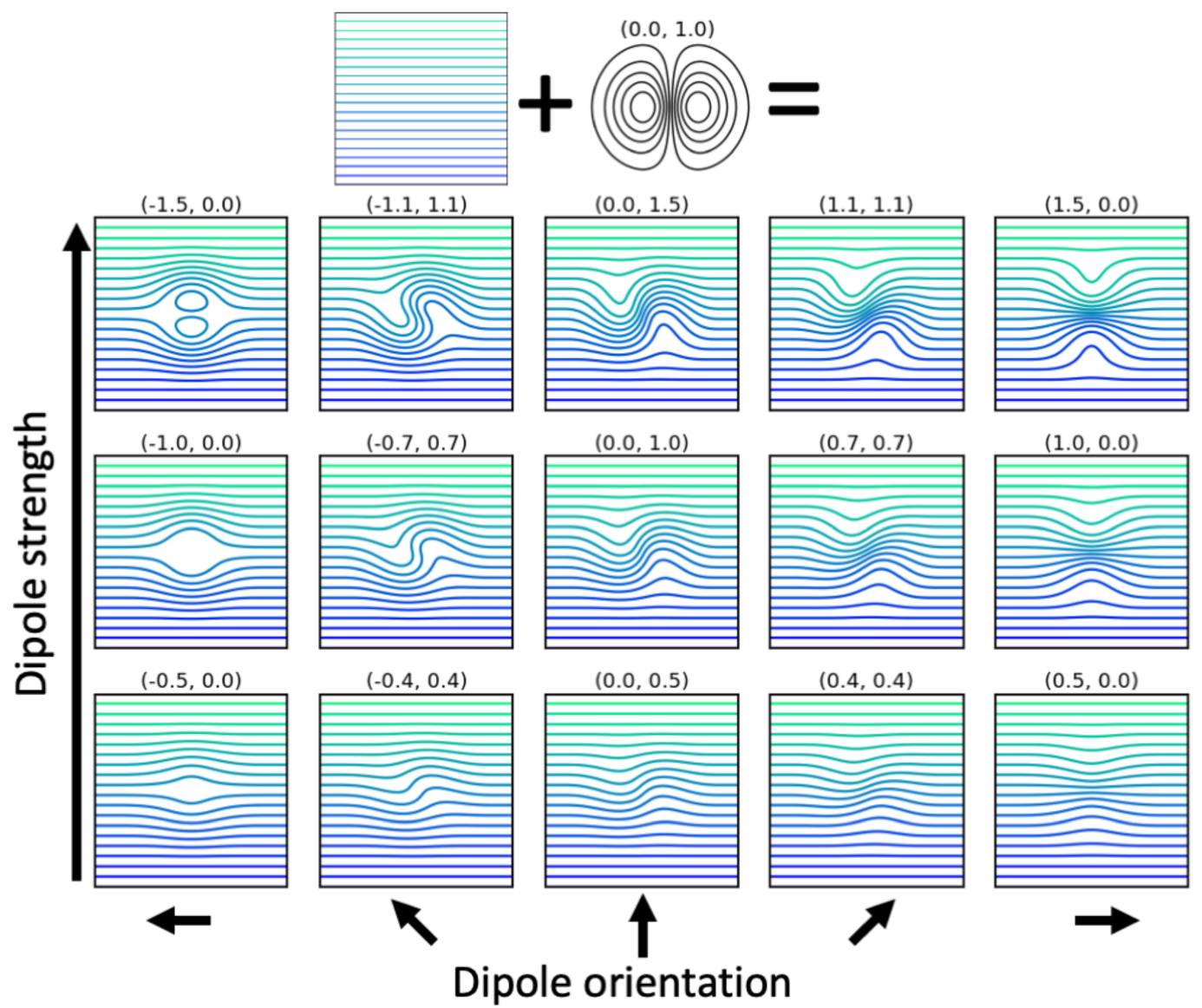
Effect of adding dipoles to a uniform vector field running from left to right. The numbers above each panel show the weights of the dipole, which set the dipole strength in respectively the x- and y-direction. Because the dipoles are divergence-free, they only alter the shape of the existing streamlines rather than allow them to terminate or reconnect. However, for sufficiently strong dipoles new, closed streamline loops might be generated (see panel in upper left).

The dipole basis functions are used to update the field distribution from the charges (eq. 2) given a set of weights 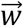. In order to efficiently evaluate the vector field 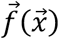 we choose to restrict ourselves to a linear and sparse mapping *M* between the parameters and the vector field:

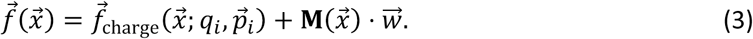

Hence the vector field is modelled as a linear combination of the columns in the matrix **M** (which represent the individual dipole-like basis functions). The shape of the matrix 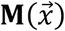 is 3x*N*, where each row defines the x-, y-, and z-component at position 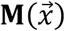 for each of the *N* basis functions. This vector field will be divergence-free by construction if each individual column in the matrix **M** is itself divergence-free, because the divergence operator is linear. The vector field generated by actual dipoles extend infinitely far from the dipole and they are therefore challenging to handle as a basis function in our application. Infinite extent of dipoles means that the mapping 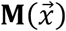 defined above will be dense, which makes the optimisation of any non-linear cost function unfeasible. Instead, we devise *dipole-like basis functions* that only extend a limited range from the dipole centre, which ensures that the mapping 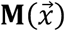 is sparse and hence the optimisation is tractable.

We start the construction of these dipole-like basis functions by defining a radial basis function. A radial basis function is any scalar function 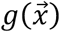 that only depends on the distance from some control point 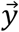. To ensure the sparsity of 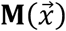 we use compactly supported radial basis functions (Wendland 1995; Buhmann 2000), which are only non-zero within a sphere around the control point, i.e. 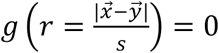 if *r* > 1, where *r* is the distance to the control point normalized by the extent of the radial basis function *s*. Here we use a compactly supported radial basis function from Wendland (1995):

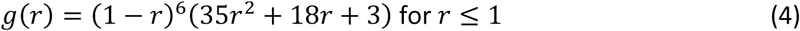

This radial basis function has the advantage that it is 4 times continuously differentiable at both *r* = 0 and *r* = 1 in 3-dimensional space (Wendland 1995), which means there will be no discontinuities in the vector field (or its first derivative), when defined from the second derivatives of this radial basis function.

The vector field basis functions are defined as 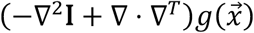 (Narcowich and Ward 1994), where 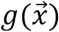 is the radial basis function defined above (eq. 4). The operator (−∇^2^ **I** + ∇ ⋅ ∇^*T*^) is chosen, so that the columns of the resulting matrix are divergence-free and hence can be used as basis functions. These columns are:

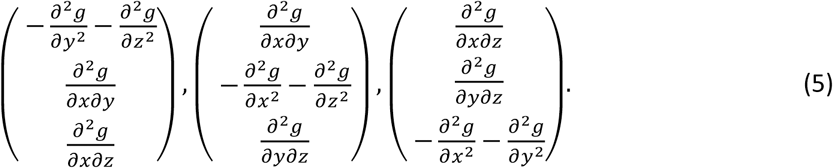

We can define *N* different radial basis functions 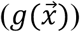 by defining these dipole-like basis functions around *N* control points. This will give us 3*N* basis functions (eq. 5), whose contribution to the vector field is determined by 3*N* parameters (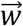; eq. 3). To fit to an arbitrary orientation field, we place these dipoles in a hexagonal grid with the distance between neighbouring dipoles given by 1/3 of the size *s* of their full extent. When these dipole-like fields are embedded within a larger field they can locally alter the shape of the field in 3 dimensions (Figure 4) to fit any target density or vector orientation, e.g. white matter orientations estimated from diffusion MRI.

The mapping from the weights 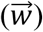 to the vector field 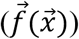 described above has been implemented in the accompanying code for both CPU and GPU. On both CPU and GPU, the matrix 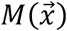 can either be pre-computed to allow for fast evaluation or can be computed on the fly if there are memory constraints.

Cost function and anatomical constraints

We use both the geometry of the cortical folds as well as fibre orientations estimated from diffusion MRI data to constrain the shape and density of our white matter model (i.e., optimise the strength and orientation of the dipoles defined in section 3.2). Here we discuss the terms adopted for the cost function in this work. Additional terms available in the accompanying code are listed in Table 1.

**Table 1.**
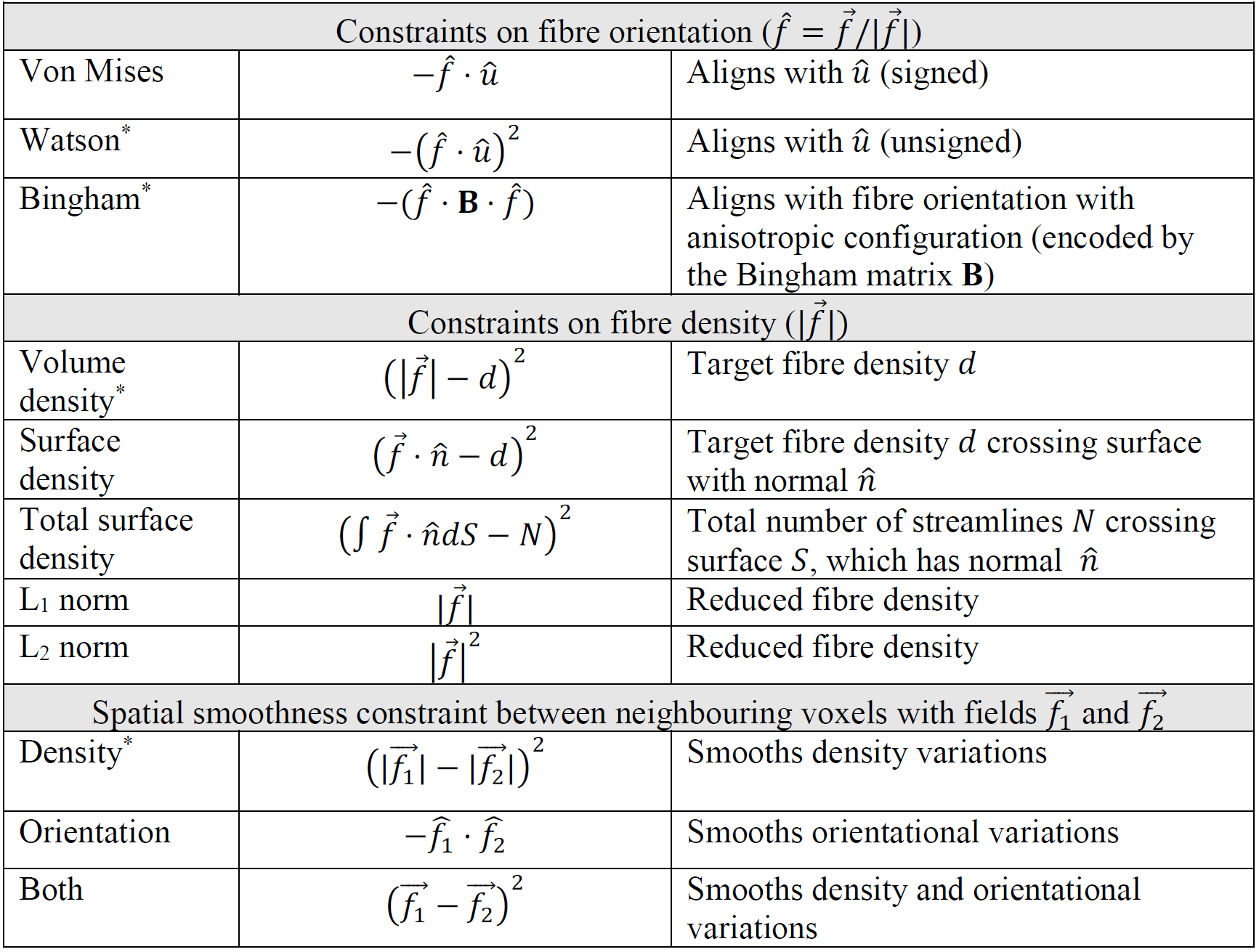
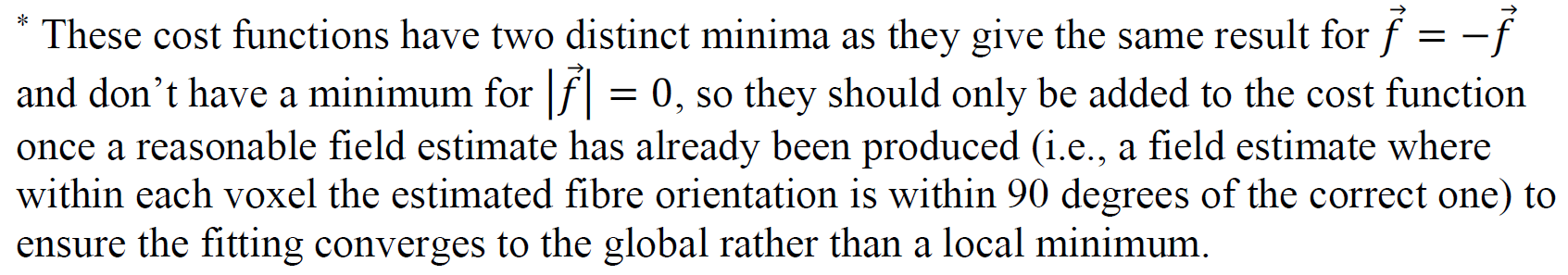
List of the available cost functions to constrain the fibre distribution.

For white matter voxels within the gyral blades our main data fidelity term in the cost function constraint will be encouraging alignment with the fibre orientations estimated from the diffusion data. As we do not model crossing fibres in this work, we define this by alignment between the principal eigenvector of the best-fit diffusion tensor 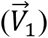 and the vector field 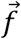 averaged over each voxel:

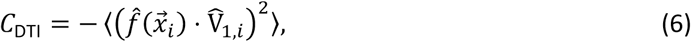

where the triangular brackets ⟨·⟩ refer to taking the average across all voxels. Note that this is a constraint on the normalized vector field 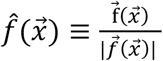, since we don’t have access to voxel-wise estimates of the fibre density in the tensor model. Because this constraint adds a degeneracy to the cost function by giving the same result for 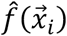 and 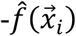, we only add it to the cost function once a decent initial estimate of the vector field has been obtained.

To encourage a smooth density distribution throughout the white matter, we also add an L2 norm constraint to the streamline density:

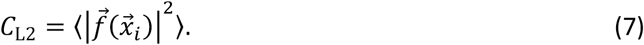

We set additional constraints at both the white/grey-matter boundary and mid-cortical surface (i.e., a mesh halfway between the white/grey-matter boundary and the pial surface). These constraints are applied to the vector field 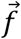 averaged over each triangle in the cortical surfaces. A constraint on density at the surface is defined for a given target density *d*_*i*_ by:

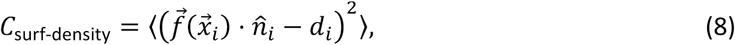

where 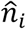 is the surface normal. The target surface density is set to encourage a uniform density of streamline endpoints through the cortical grey matter volume (Van Essen et al. 2014).

Finally, a radial fibre orientation at the surface is encouraged by including the following term to the cost function:

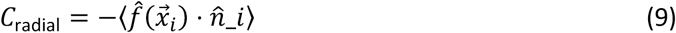

Minimising this term will maximise the alignment between the orientation of the vector field and the surface normal. The total cost function is then given by:

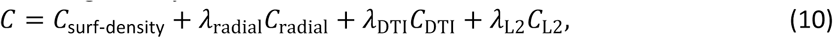

where the individual cost functions are defined in eqs. 6-9 and the *λ*s give the relative weights of the different cost functions (which will be given in section 3.5).

### 3.4 Interface with probabilistic tractography

The vector field produced by optimising the cost function above can be thought of as providing a one-to-one mapping between any location on the cortical surface with a location for streamlines to enter into deep white matter. We take each vertex on the cortical surface and move it along the vector field as described below to the interface between the gyral and deep white matter (right in Figure 2). This creates a deformed but topologically equivalent version of surface around deep white matter, which excludes the cortical convolutions. This surface can then be used as a seed and/or target mask in any tractography algorithm.

### 3.5 Building whole-brain connectomes

Given cortical surface models, surfaces (e.g. extracted from an anatomical T1w image) and diffusion MRI data for a single subject, we build a *dense* (i.e. vertex/voxel-wise rather than parcel-wise) connectome using the following steps:

- First, we create a mask of the white matter within the gyral blades. This mask is designed to include any voxels within the gyral white matter and includes those voxels through which the shortest line connecting the gyral walls on both sides is shorter than 10 mm (Figure 2).
- Within the gyral white matter we estimate the fibre configuration in three steps:
  1. An initial estimate of the vector field is generated by placing uniform “negative” charges across the pial surface. These charges are compensated for by an equal-sized positive charge in the centre of the deep white matter in each hemisphere. These charges generate a vector field flowing from the pial surface into the brain according to eq. 2.
  2. This initial field is refined using the dipole-like basis functions (eqs. 3-5). These basis functions have a finite extent of *s* (=20 mm used here) and are interspersed on a hexagonal close packing configuration at a distance of ^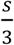^ mm. The strength and orientation of the dipoles is optimised by minimizing the cost function using the quasi-Newton method L-BFGS-B (Byrd et al. 1995; Zhu et al. 1997):

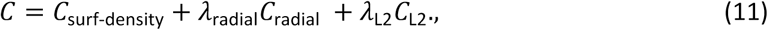

which encourages a uniform density of streamline endpoints in the cortical volume (eq. 8), a radial orientation at the cortical surface (eq. 9) and imposes an L2 norm on the volumetric fibre density (eq. 7). Both surface constraints Csurf-density and Cradial are enforced at the white/grey-matter boundary as well as the mid-cortical surface.
  3. Finally, a set of smaller dipoles (extent of 7 mm, interspersed on a hexagonal grid with distance of 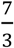 mm) is optimized by the cost function above and additionally enforcing alignment between the principal eigenvector of the diffusion tensor and the proposed fibre orientation (eq., 6)

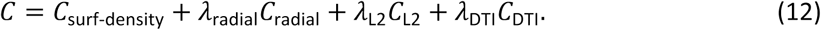 Here we use *λ*_radial_ = 1, *λ*_DTI_ = 1, and *λ*_L2_ = 10^−3^. These values were determined through trial-and-error based on the quality of the resulting fit and visual inspection of the resulting vector field. How these parameters should be set to robustly work across a large number of datasets, remains to be investigated. The final vector field is given by the sum of the contribution of the initial field (step 1), the large dipoles (step 2) and the smaller dipoles (step 3).
- The final vector field is used to guide the vertices of the white/grey-matter boundary through the gyral white matter. This provides a 1:1 mapping and results in a new deformed surface at the interface between the gyral and deep white matter that encloses deep white matter.
- During tracking from the white/grey-matter boundary neighbouring vertices do not always remain immediately adjacent to one another other, which leads to a very ragged-looking mesh. To resolve this, we smooth the mesh at the gyral and deep white matter interface by moving each vertex towards the mean of its neighbours. During this smoothing the vertices moved a median distance of less than 1 mm, with 95% of vertices moving less than 3 mm. The smoothed surface (which has a mesh density that varies greatly across the interface) was used as seed and target for tractography.

### 3.6 Data and analysis

We tested our algorithm on pre-processed data from 20 subjects of the Human Connectome Project (HCP) (Van Essen et al. 2012). The pre-processed data includes white/grey-matter boundaries and pial surfaces extracted from the T1-weighted and T2-weighted images using the HCP Pipelines (Glasser et al. 2013). The diffusion constraint was obtained by fitting a diffusion tensor to the b=1000 shell of the pre-processed HCP diffusion MRI data (Sotiropoulos et al. 2013; Andersson and Sotiropoulos 2016). Group-average analysis were carried out on datasets aligned using MSMAll intersubject registration (Robinson et al. 2014; Glasser et al. 2016).

We used FSL’s probtrackx2 (Behrens et al. 2007; Hernandez-Fernandez et al. 2019) to compare the features of the connectome when seeding/terminating streamlines at either the white/grey-matter boundary or at the new interface between the gyral and deep white matter. For these two surfaces we (i) compared the density distribution of streamline endpoints when seeding from the subcortical volume or from the contralateral hemisphere, (ii) assessed the similarity in the path that streamlines seeded from the surface take through deep white matter, and (iii) performed a comparison between the functional and structural connectome. In each case the structural connectivity was estimated by dividing the number of streamlines connecting two voxels or vertices by the cortical volume associated with the target vertex or voxel. This makes our structural connectivity from A to B measure proportional to the probability of streamlines seeded in voxel/vertex A to terminate in each mm^3^ of voxel/vertex B.

In four subjects these connectomes were parcellated using the subject-specific multi-modal parcellations from Glasser et al. (2016). The connectivity from parcel A to parcel B was estimated by adding up all the streamlines going from A to B and then divide by the number of vertices in A and the total cortical volume associated with B. This connectivity measure is once again proportional to the average probability of streamlines seeded in parcel A to terminate in each mm^3^ of parcel B. This parcellation allows quantification of the connectivity between homotopic and heterotopic interhemispheric connections.

For comparison with tracer data from Markov et al. (2011; 2014) we also compute a parcellated connectome in an ex-vivo macaque diffusion MRI dataset in the same manner as described above, except for scaling down the threshold to define gyral white matter (10 to 4 mm) and the size of the dipoles (in initial fit from 20 to 9 mm, in final fit from 7 to 3 mm) to compensate for the smaller brain size. As the tracer data is reported as the fraction of labelled cells within a given ROI, we also apply such fractional scaling to the connectome from tractography using the algorithm described by Donahue et al. (2016). The diffusion MRI data and its preprocessing have been previously described in Jbabdi et al. (2013). In summary ex-vivo diffusion data was acquired on a 4.7 T scanner using a 3D-segmented spin-echo EPI sequence (430 μm isotropic resolution, TE=33 ms, TR=350 ms, 120 directions, *b*_max_= 8000 s/mm^2^).

Results that are displayed as Connectome Workbench scenes are available via the BALSA database (https://balsa.wustl.edu/study/show/0LGM2). Code, documentation, and a tutorial of the proposed algorithm can be found at https://git.fmrib.ox.ac.uk/ndcn0236/gyral_structure.

## 4 Results

First, we defined for each subject a gyral white matter mask including those white matter voxels that lie between the gyral folds (Figure 2). Within this gyral white matter we found the best-fit vector field (by minimising eq. 10) that aligns with the primary eigenvector of the diffusion tensor and is both uniform and radial at the white/grey-matter boundary and mid-cortical surface.

Figure 5 shows maps of the best-fit vector field density and orientational alignment with the diffusion tensor for a sample subject as well as histograms of the full distribution for both hemispheres in 20 subjects. Consistently across both hemispheres in 20 subjects we find an excellent alignment with the diffusion tensor primary eigenvector (Figure 5B) in all regions apart from the boundaries where the vector field becomes radial, as well as a fairly uniform density distribution at both the white/grey-matter boundary (Figure 5C) and the mid-cortical surface (Figure 5E). While the orientation field has become mostly radial at the mid-cortical surface (Figure 5F), at the white/grey-matter boundary the field is still far from radial for large parts of the cortical surface (Figure 5D). The radiality can be improved by increasing its influence in the cost function or reducing the size of the dipoles in the basis function (which allows for sharper curvature of the vector field), however the lack of perfect radiality at the white/grey-matter boundary is expected in a realistic fibre configuration (Budde and Annese 2013; Reveley et al. 2015; Cottaar et al. 2018).

**Figure 5.**
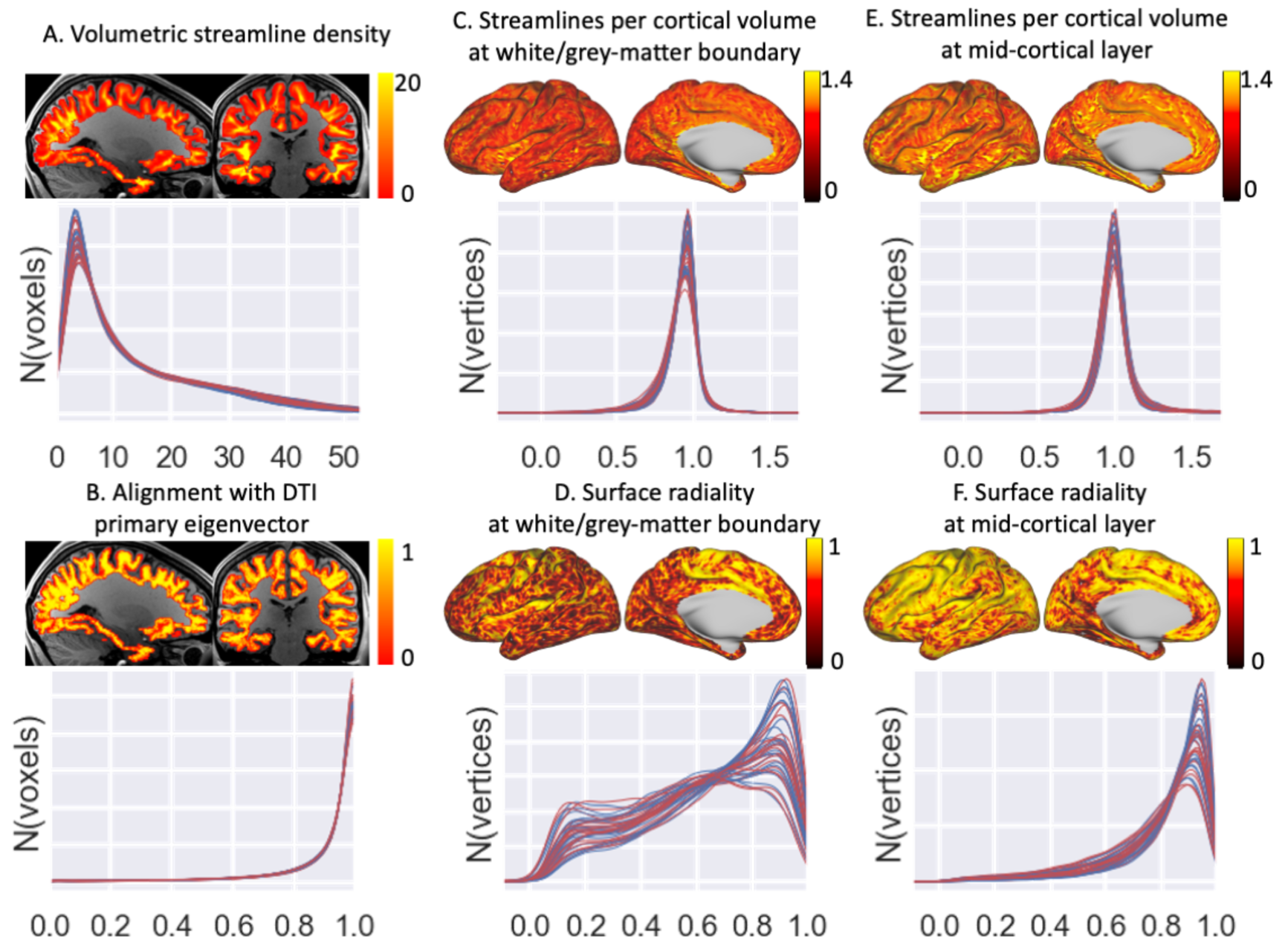
The distribution of the different terms of the cost function for the best-fit vector field. This cost function includes an L2 norm on the volumetric streamlines density (A), increases alignment with DTI V1 (B), approximates a uniform density per cortical volume element across both the white/grey-matter boundary (C) and the mid-cortical surface (i.e., halfway between the white/grey-matter boundary and the pial surface) (E), and finally increases alignment with the surface normal at both surfaces (D & F). For each variable a volumetric or surface map is shown for a single subject and the density distributions for 20 subjects (left hemisphere in blue and right hemisphere in red). Note that this plot illustrates the density of the best-fit vector field in the superficial white matter. The density of this vector field might not reflect the density of streamlines resulting from tractography running through the deep white matter (which is illustrated in later figures).

While the L2 norm (eq. 7) attempts to reduce the volumetric streamline density (Figure 5A), the divergence-free constraint limits its effectiveness as the streamlines crossing the surface must go somewhere. The weight on the L2 norm is chosen to be low enough not to significantly lower the number of streamlines crossing the white/grey-matter boundary, but high enough that it discourages those streamlines from taking a circuitous route through the gyral white matter (which would increase the average streamline density).

The resulting best-fit fibre configuration is illustrated in Figure 6 for a few gyri. This vector field is used to guide the vertices from the white/grey-matter boundary to the deep white matter. This creates a new deep/gyral white matter interface (blue) where each vertex has a one-to-one correspondence with the white/grey-matter boundary (turquoise). Note that the deep/gyral white matter interface shown here has been smoothed.

**Figure 6.**
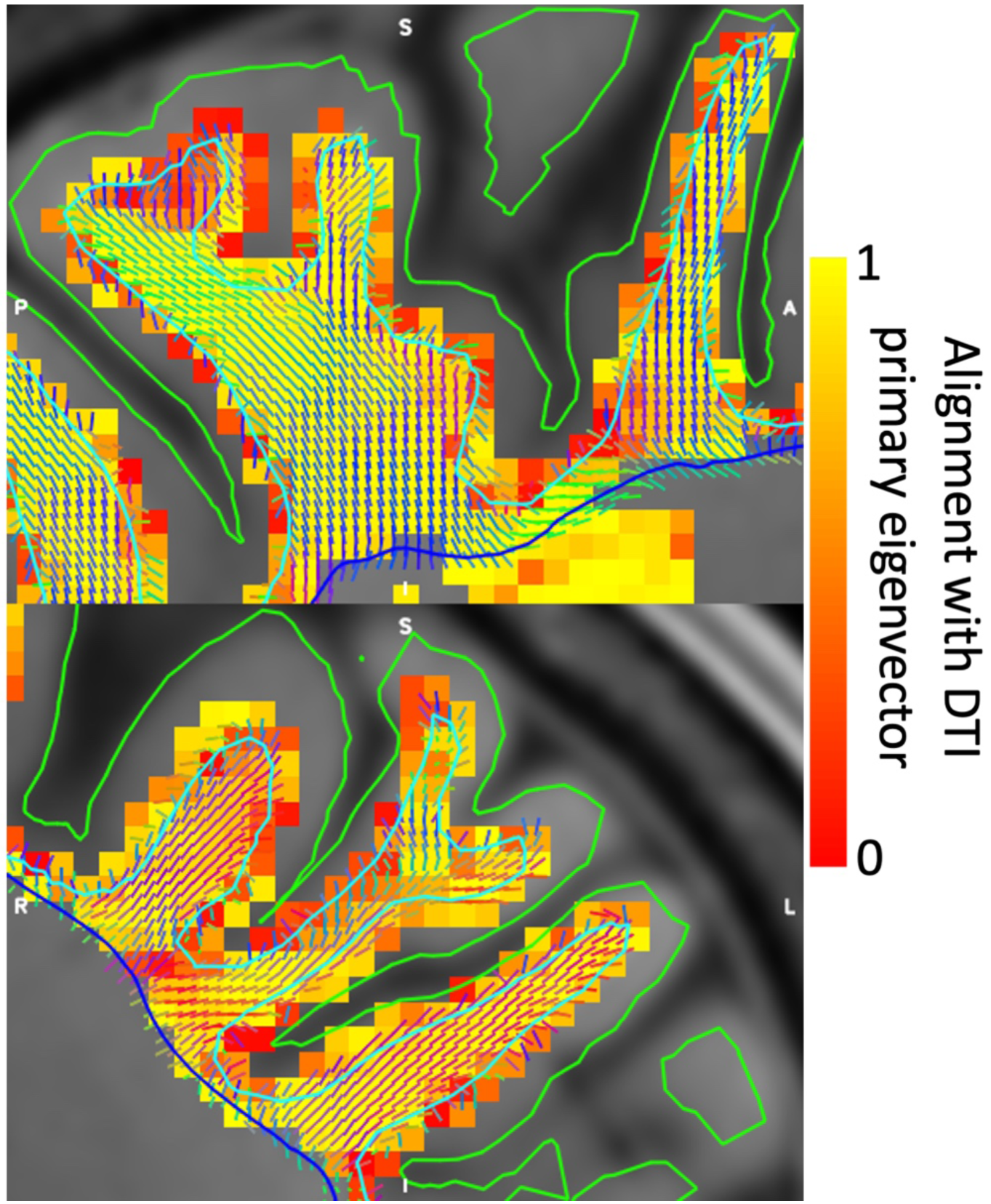
Vector field configuration in sample gyri for a single subject. Note that the vector field itself is a continuous 3D function defined at every intermediate point, but here we discretise it by averaging the vectors within each image voxel and showing a grid of these mean vectors extracted from the vector field. The colour map shows the absolute value of the dot-product between the continuous vector field sampled at the centre of each voxel and the primary eigenvector of the diffusion tensor at that voxel. The deep/gyral white matter interface (blue) has a one-to-one vertex correspondence with the white/grey-matter boundary (turquoise) and pial surface (green).

Figure 7 compares the density of the estimated vector field (A) with the streamline density from seeding tractography at the white/grey-matter boundary (B) or the deep/gyral white matter interface (C). While seeding from the white/grey-matter boundary is (by construction) uniform on the surface, the resulting distribution is very non-uniform in the gyral white matter (left in Figure 7B). Streamlines tend to stick closely to the white/grey-matter boundary following the U-fibres and relatively few reach deep white matter. On the other hand, streamlines seeded from deep/gyral white matter interface tend to have a higher density in the central part of the gyri and avoid the white/grey-matter boundary (left in Figure 7C) until they reach the top of the gyral crown (right in Figure 7C). Our vector field model uniformly connects the white/grey-matter boundary with most (although still not all) of the gyral white matter (Figure 7A).

**Figure 7.**
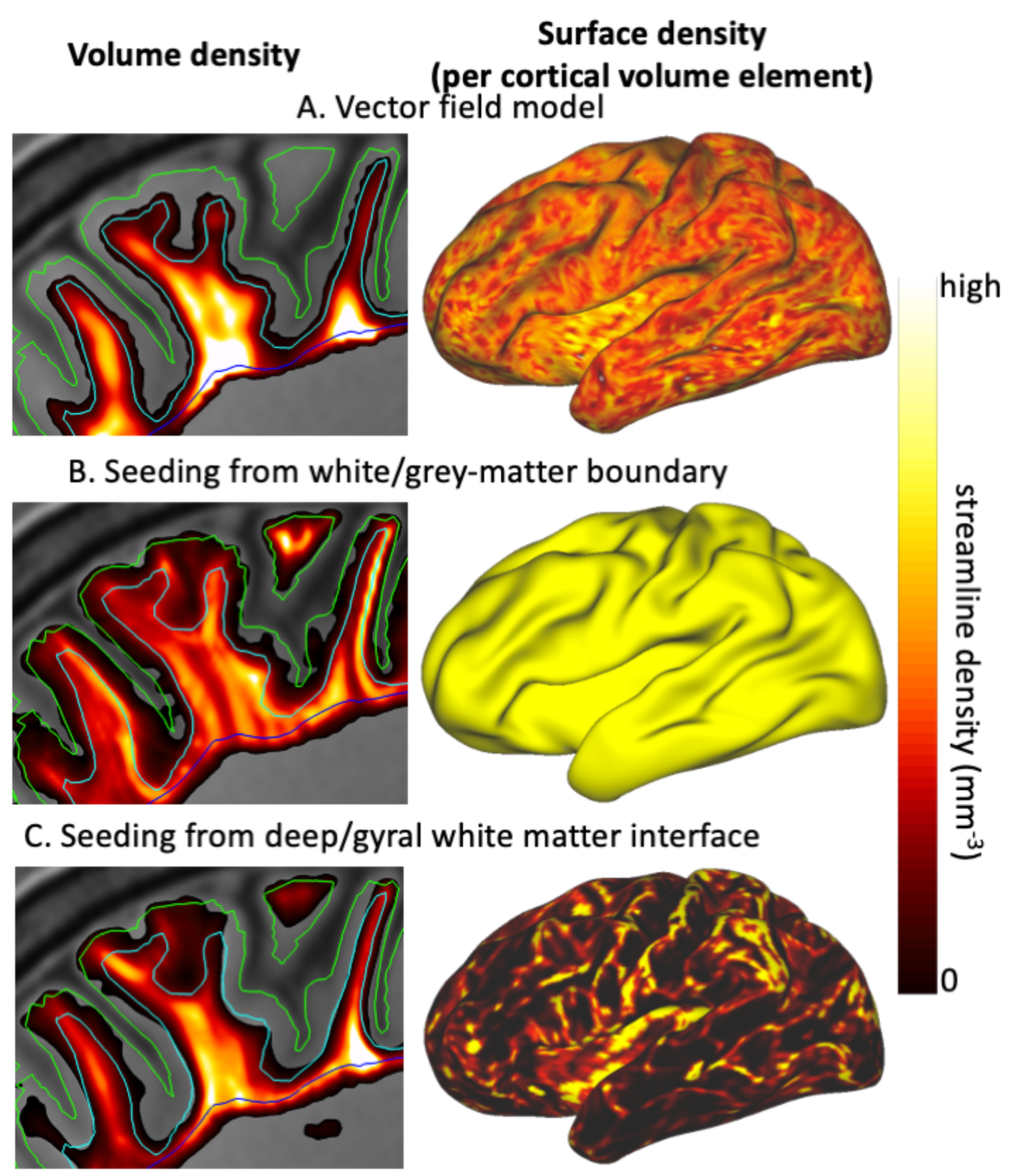
For a single subject volumetric density (left) and surface density (per cortical volume element) at the white/grey-matter boundary (right) for the vector field model (A), probabilistic tractography from the white/grey-matter boundary (B) and from the deep/gyral white matter interface (C). Overlaid are the pial surface (green), white/grey-matter boundary (cyan) and deep/gyral white matter interface (blue). Because the streamline density has very different scaling in the different panels, the density in each panel was normalised independently before applying the same linear mapping to colour. While the vector field has a smooth density in the white matter (A; left) and on the surface (A; right), tractography seeded from the white/grey-matter boundary leads to a bias of streamlines close to the cortex (B; left), while tractography seeded from the deep/gyral white matter interface has a strong gyral bias on the surface (C; right).

It is worth noting that even if the vector field describing the gyral white matter is uniform per cortical volume element, this does not guarantee that the tractography streamlines will be uniformly distributed per cortical volume element after travelling through deep white matter. The vector field merely provides a one-to-one mapping between points on the cortical surface and points at the interface between the deep and gyral white matter. Whether this leads to a reduction in the gyral bias depends on the distribution of streamlines along this deep/gyral white matter interface.

Thus, to further investigate the gyral bias, we run tractography streamlines seeded in the contralateral cortex and subcortical grey matter regions (as defined in the HCP grayordinate space) (Glasser et al. 2013) up to either the deep/gyral white matter interface (top in Figure 8) or the white/grey-matter boundary (bottom in Figure 8). Due to the one-to-one correspondence of the vertices between the two surfaces, we can assign each streamline terminating at the deep/gyral white matter interface to the equivalent vertex at the white/grey-matter boundary. This is equivalent to propagating these streamlines to the white/grey-matter boundary along the best-fit vector field.

**Figure 8.**
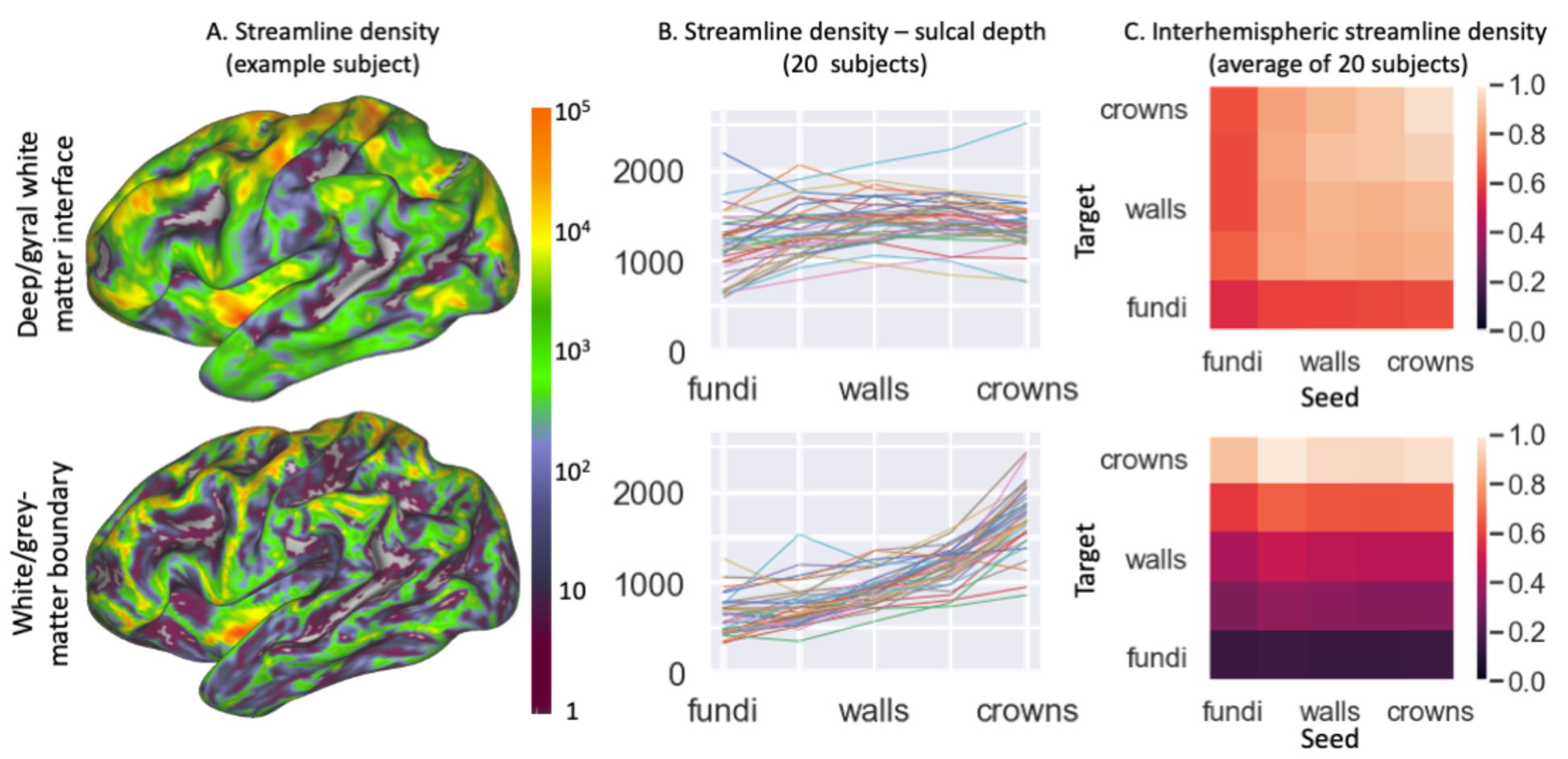
Reduction in gyral bias by tracking to the deep/gyral white matter interface (top) rather than to the white/grey-matter boundary (bottom). Seeding is from subcortical grey matter and the contralateral cortex. A) Streamline termination density per mm^3^ of cortex on left cortical surface for a single subject using a logarithmic scale spanning five orders of magnitude. B) Streamline termination density per mm^3^ of cortex for five sulcal depth bins (all bins have an equal total area on the mid-cortical surface). C) Streamline termination density per mm^3^ of one hemisphere per 10^6^ streamlines seeded in the contralateral hemisphere for the same five sulcal depth bins.

When only considering these streamlines from other grey matter brain regions, the large effect of the gyral bias can be appreciated. Tens of thousands of streamlines terminate in part of the cortex (in particular the gyral crowns and the insula), while large parts of the cortex get no streamlines at all (bottom in Figure 8A). When terminating at the deep/gyral white matter interface a great increase in the coverage can be seen (top in Figure 8A), however many of the sulcal fundi are still not covered (see Figure S3A for a similar result in the macaque). This corresponds to a reduction in the dependence of the streamline density on sulcal depth (Figure 8B).

Figure 8C illustrates in more detail the connectivity profile of the commissural streamlines. Commissural streamlines seeded at the white/grey matter surface are very likely to terminate in the gyral crown of the contralateral white/grey matter surface (bottom in Figure 8C). While this trend is reduced for the deep/gyral white matter interface, some preference for terminating at the gyral crown in still present (top in Figure 8C). The same preference for gyral crowns is now found for streamline traveling in the other direction, with streamlines seeded from the gyral crowns being more likely to reach the contralateral cortex (top in Figure 8C). It is unclear whether this remaining dependence on sulcal depth is genuine, but in any event its magnitude is minor compared with the gyral bias observed when tracking between the contra-lateral white/grey matter boundaries (bottom in Figure 8C).

Next, we investigate the behaviour of streamlines seeded from the cortical surface, rather than the gyral bias of those approaching the surface. Figure 9 illustrates the dissimilarity of the path streamlines take through deep white matter between neighbouring vertices. A large dissimilarity corresponds to a sudden change in the structural connectivity profile, indicating a potential border between two distinct cortical areas (Johansen-Berg et al. 2004; Fan et al. 2016).

**Figure 9.**
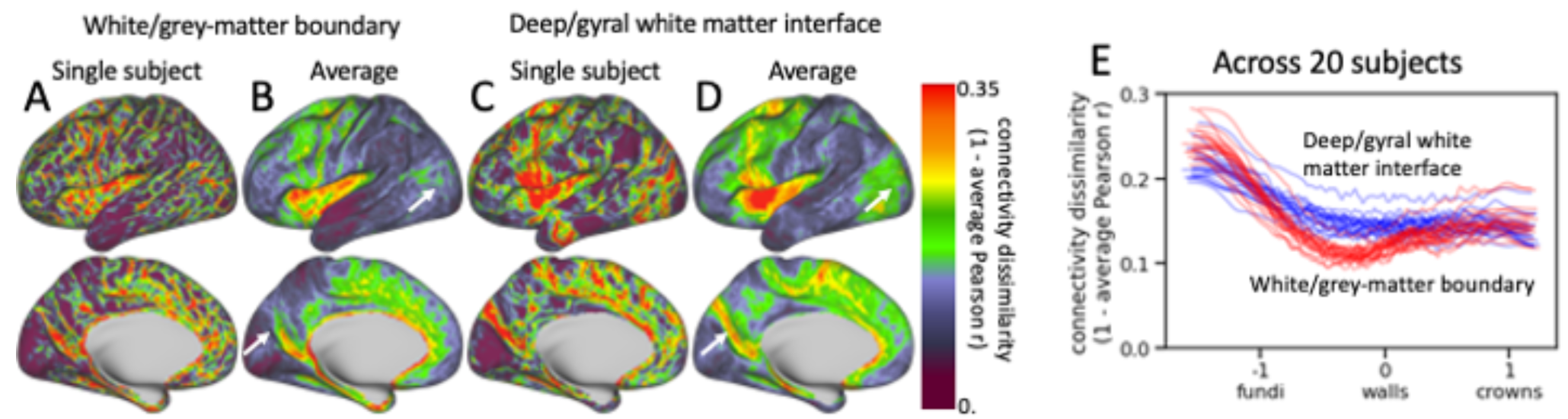
Dissimilarity of the structural connectivity profiles in the deep white matter between neighbouring vertices for streamlines seeded from the white/grey matter surface or the deep/gyral WM interface. The dissimilarity is computed as one minus the Pearson-r correlation across the connectivity with all voxels below the deep/gyral WM interface. High dissimilarity indicates that streamlines seeded from that vertex take a very different path through the deep white matter from the neighbouring vertices (i.e., there is a strong gradient in structural connectivity). Top: dissimilarity maps for a single subject (A & C) and averaged across 20 subjects (B & D) for streamlines seeded from the white/grey-matter boundary (A & B) and the deep/gyral white matter interface (C & D). White arrows point to the parieto-occipital sulcus; E: trend lines of the dissimilarity with sulcal depth for 20 subjects (each line represents a single subject) with seeding from the white/grey-matter boundary in red and seeding from the deep/gyral white matter interface in blue. Trend lines were created using median-filtering of the dissimilarity across 400 vertices after sorting by sulcal depth.

When seeding from the white/grey-matter boundary, narrow strips with high dissimilarity are widespread across the cortex (Figure 9A). These tend to follow the gyral crowns and sulcal fundi with streamlines seeded from the gyral walls being very similar between neighbouring vertices (Figure 9E). This likely reflects the tendency of streamlines seeded from the gyral walls to stick close to the cortex as illustrated in Figure 7B, which causes streamlines seeded from the gyral walls to enter the deep white matter close to each other. When seeding from the deep/gyral white matter interface this alignment of the structural connectivity gradient with the gyrification is reduced (Figure 9C), although on average the dissimilarity remains largest in the sulcal fundi (Figure 9E).

When averaging across subjects, most of the detail in these structural connectivity boundary maps disappears (Figure 9B,D). Still, some plausible boundaries remain such as at the edge of the occipital lobe (marked by white arrows), particularly on the medial side in the parieto-occipital sulcus. These boundaries are less well defined when seeding from the white/grey-matter boundary than from the deep/gyral white matter interface, which likely reflects the better alignment of the structural connectivity profile gradients when the effect of the gyrification on the tractography is reduced.

Figure 10 compares the estimated group structural connectivity profiles for selected seeds when using the deep/gyral white matter interface rather than the white/grey-matter boundary. For comparison the average functional connectome across all subjects as downloaded from the HCP database has been included on the left. While the structural connectivity profiles are generally similar, many differences are evident. In general, there appear to be more long-distance connections when seeding from and targeting the deep/gyral white matter interface. In some regions this improves the agreement with the functional connectome (green arrows in Figure 10), although counter-examples can also be found (purple arrow in Figure 10).

**Figure 10.**
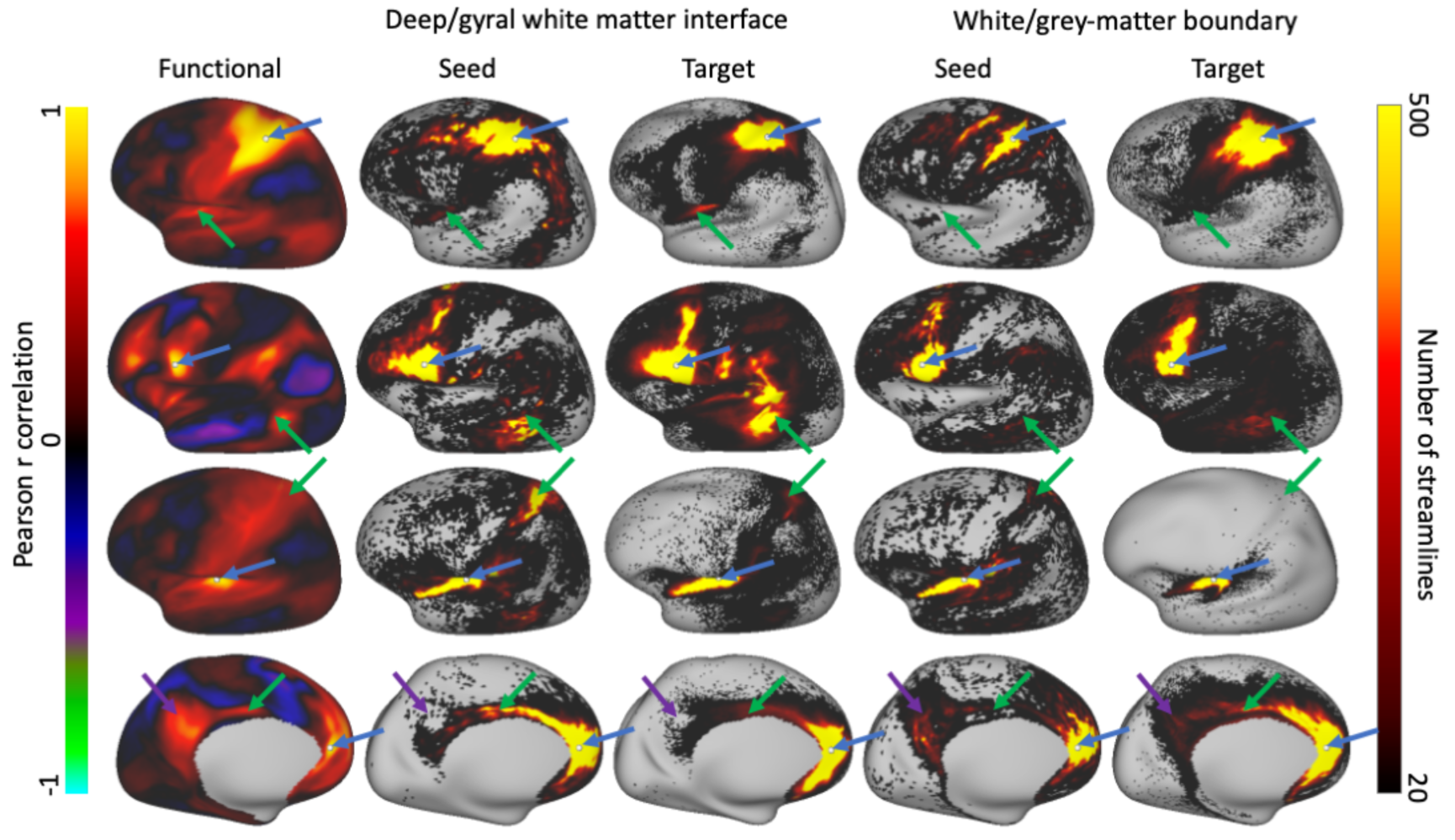
Comparison of the structural connectome (averaged over 20 subjects) with the functional connectome (averaged over all HCP subjects). The connectivity from a reference vertex with from left to right: functional connectivity, structural connectivity when using deep/gyral white matter interface (using reference vertex as seed on left or as target on right), and finally structural connectivity using white/grey-matter boundary (again using reference vertex as seed on left or as target on right). From top to bottom reference vertices are in the parietal lobe, frontal lobe, insula, and cingulate (marked by white dots and the blue arrow). Green arrows mark distant intrahemispheric connections where the agreement with the functional connectome seems to have improved when using the deep/gyral white matter interface, while the purple arrow marks an area where using the white/grey-matter boundary works better.

To quantify the comparisons between these 3 connectomes (i.e., streamlines seeded from a reference vertex, streamlines terminating in a reference vertex, and the functional connectivity) we compute the Pearson correlation between them for every vertex (Figure 11). Overall, the correlations between the (log-transformed) structural and functional connectome are very low (left two columns), whether we consider nonlocal intrahemispheric connections (top), interhemispheric connections (middle) or connections with the subcortex (bottom). A slight improvement in the correlation is seen in the interhemispheric connections when adopting the deep/gyral white matter interface. Adopting the deep/gyral white matter interface does greatly boost the symmetry of tractography, with the distribution of streamlines seeded from a vertex being more similar to the distribution of streamlines terminating in a vertex (right column in Figure 11).

**Figure 11.**
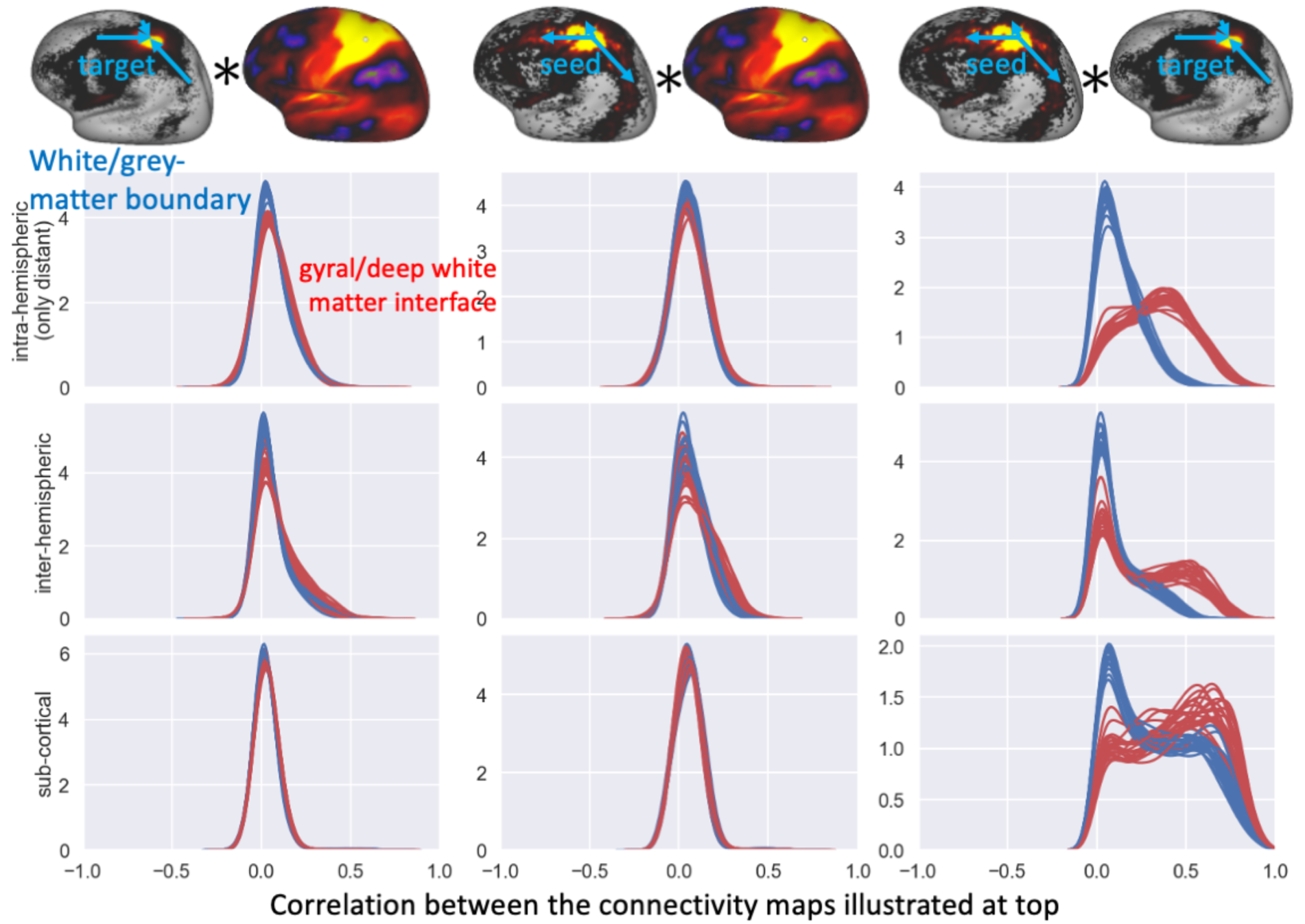
Distribution of the correlations between the structural and functional connectivity profiles when seeding from the white/grey-matter boundary (blue) or gyral/deep white matter interface (red) with each line showing the distribution for one out of 20 subjects. For each vertex the correlation is computed between the connectivity estimates with respect to either all other vertices in the same hemisphere excluding local and U-fibres as defined in Figure S1 (top panels), or all vertices on the contralateral hemisphere (middle panels), or all sub-cortical grey meter voxels as defined in the HCP grayordinate space (bottom panels). As illustrated at the top, the correlations are computed between the functional connectome with either the log-density of streamlines terminating in a vertex (left panels) or the log-density of streamlines seeded in a vertex (centre panels). The right panels compare the log-density of the two structural connectivity profiles (i.e., seeding from or targeting a vertex).

So far, we have exclusively focussed on the dense (i.e., vertex-wise) connectome. Further validation can be obtained by studying the parcellated connectome. We use the multi-modal parcellation from Glasser et al. (2016) to parcellate the cortical connectome (Figure S2A). Because we only alter the tractography within the gyral blades, the connectivity strengths in these parcellated connectomes are strongly conserved between using the white/grey-matter boundary or deep/gyral white matter interface (Figure S2B,C). However, these minor changes in the parcellated connectomes still allow for some additional validation. The results of this experiment appear to be mixed. In the HCP data adopting the deep/gyral white matter interface increases the interhemisphere connectivity between homotopic regions, while decreasing the interhemispheric connectivity between heterotopic regions (Figure S2D), which is in line with the predominance of homotopic connections seen in tracer studies (e.g., Oh et al. 2014). However, when applied in a macaque diffusion MRI dataset previously described in Jbabdi et al. (2013), the correlation with the “ground-truth” connectome based on neuroanatomical tracers from Markov et al. (2011; 2014) decreases.

## 5 Discussion

Here we present a model for the white matter in gyral blades, which reduces the overestimation of gyral connectivity and underestimation of sulcal connectivity by considering the shape of the gyrus when running tractography in the gyral white matter (Figure 8). This is done by imposing two physical constraints on the gyral white matter fibre configuration: (1) fibres do not terminate in the white matter (i.e., the vector field is divergence-free) and (2) fibres do not cross each other. The first continuity constraint ensures that all these streamlines uniformly entering the gyral white matter have to go somewhere and the only possible destination is deep white matter. The second non-crossing constraint ensures that when the streamlines converge on the interface with the deep white matter, those from the left gyral wall remain on the left, those from the right gyral wall remain on the right, while those from the gyral crown get compressed into the centre of the gyral white matter (Figure 1). It has previously been argued that such an assumption of spatial organisation within a white matter bundle is crucial for tractography to be able to claim any relation between where fibres enter and leave a white matter bundle (Jbabdi et al. 2015). With these constraints, we optimise a cost function to create a uniform (and radial) fibre distribution at the white/grey-matter boundary and mid-cortical surface and to align with the primary eigenvector of the diffusion tensor in each voxel. The optimisation routine is consistently able to achieve a fairly uniform distribution with excellent alignment with the DTI across all 20 HCP subjects tested here (Figure 5). While this does lead to a realistic-looking fanning fibre configuration (e.g., compare Figure 6 with (Heidemann et al. 2012; Budde and Annese 2013; Van Essen et al. 2014; Sotiropoulos et al. 2016)), this model does have some limitations.

### 5.1 Model assumptions and limitations

The method assumes that there is a one-to-one mapping from each point on the cortical surface to where the fibres enter deep white matter. There is evidence for such organisation from tracer studies, at least for long-distance fibres, such as those connecting with many sub-cortical regions and the contralateral hemisphere. Many long-distance axons (in particular those connecting to the striatum, corpus callosum, cingulum bundle or the capsules) tend to be well-clustered in a narrow “stalk” while travelling through the gyral white matter and only disperse in deep white matter (Figure 12) (Krieg 1973; Safadi et al. 2018). Hence, these long-distance fibres might be well represented by the one-to-one mapping provided by the proposed model. On the other hand, the vector field does not represent the U-fibres or other short-distance fibres. These are unlikely to follow this path to deep white matter and are found to be in general far more spread out (Figure 12). Although these fibres could be included in the model by superimposing a second (or even third) vector field on top of the single one modelled here, the fact that they are spread out suggests that they might be better represented by a model that allows for fibres to cross within a single white matter bundle, such as local probabilistic tractography or the spin-glass model by Reisert et al. (2011).

**Figure 12.**
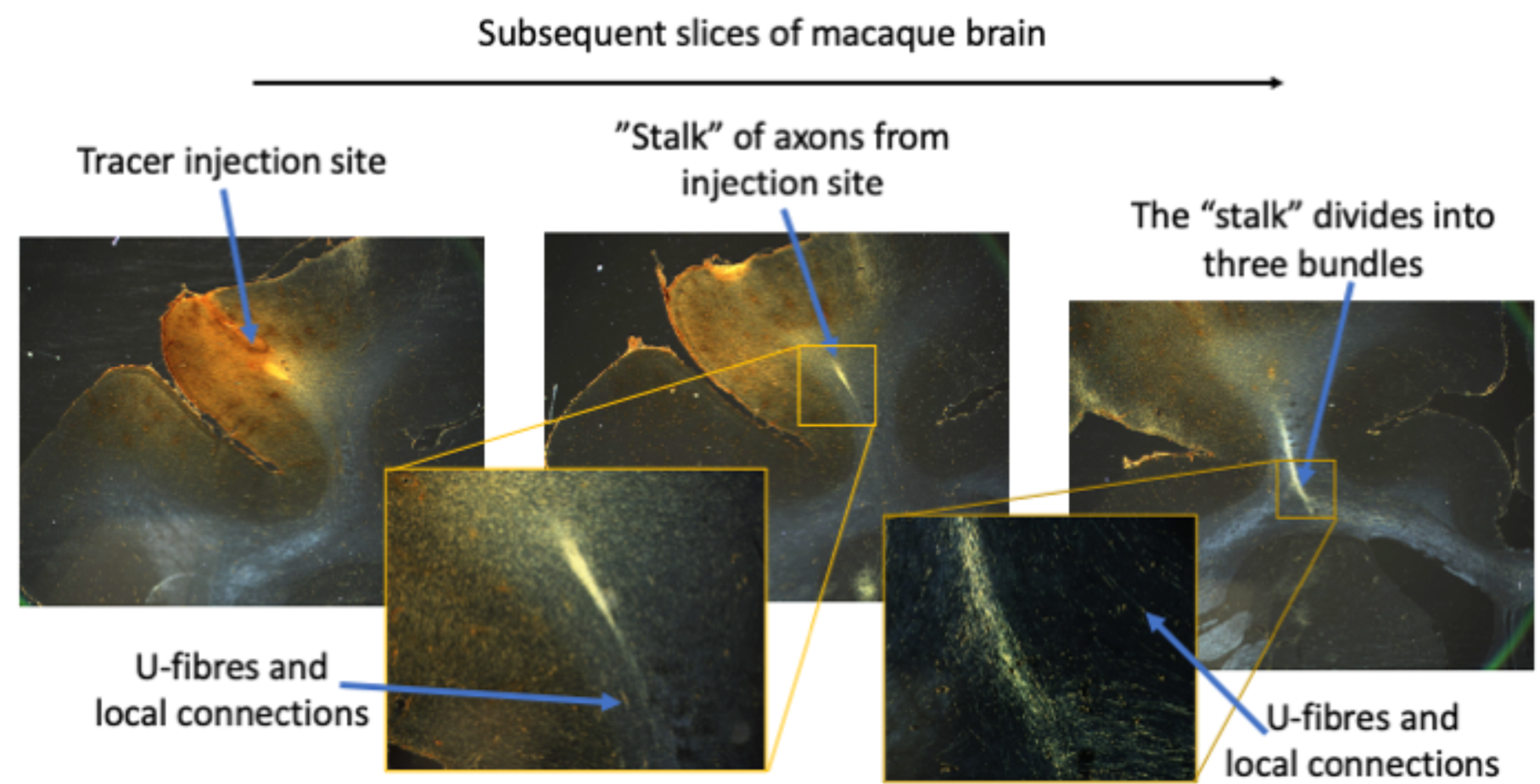
Traced axons from a bidirectional tracer (Lucifer Yellow) in the prefrontal cortex (left) of an adult male monkey (*Macaca fascicularis*). Long-distance axons can be seen to travel together from the injection site in a relatively “narrow” stalk (middle) until enter the deep white matter (right) and divide into separate bundles that travel to the corpus callosum, cingulum bundle and capsules, and the striatum. U-fibres and axons connecting within the same gyrus and those traveling to other cortical regions do not form part of this “stalk” and are far more spread out (insets). For experimental details see Lehman et al. (2011) and Safadi et al. (2018). The tracing experiment was performed in accordance with the Institute of Laboratory Animal Resources Guide for the Care and Use of Laboratory Animals and approved by the University Committee on Animal Resources at University of Rochester.

A major assumption made by the vector field model is that the fibres represented by the vector field do not cross each other. This assumption is intrinsic to our choice of modelling the fibre configuration as a vector field, where at any point we only have a single fibre orientation. Although a crossing fibre bundle could be added to the model by representing it with a second vector field (e.g., to model the U-fibres), the vector field model would still ensure that within each fibre bundle the fibres cannot cross each other. In other words, we assume that while the “stalks” seen in Figure 12 might cross the U-fibres or local axons, they do not cross “stalks” connected with different parts of the cortex (i.e., “stalks” from the left gyral wall stay on the left, those from the right gyral wall stay on the right). As far as we are aware, this assumption is as yet untested.

Finally, the target density distribution adopted in this work (i.e., a uniform streamline termination density per unit of cortical volume) is only a first-order approximation of the true expected density distribution. In reality there will be significant variation between cortical regions in the density of long-distance connections. Given the limitations of tractography in estimating the density of long-distance connections, more accurate estimates of the expected density distribution across the surface likely have to come from detailed histological studies, which is beyond the scope of this article.

### 5.2 Validation

Adopting the vector field model for the gyral white matter can be viewed as a regularisation algorithm, where we take some of the streamlines which would have terminated on the gyral crown and move them to the sulcal walls or fundi, following anatomical constraints. We show that this reduces the gyral bias when streamlines travel up to the cortex (Figure 8). By allowing streamlines not to have to track through the gyral white matter, we find many more streamlines connecting to the cortex. Still some more subtle trends with the sulcal depth remain, with commissural streamlines showing a residual gyral bias, although this bias is now the same for the hemisphere where we are seeding from and the target hemisphere (Figure 8C).

Even when seeding from the white/grey-matter boundary this reduction of the gyral bias becomes obvious when examining boundaries in the cortical connectivity profile to the deep white matter is (Figure 9). When seeding from the white/grey-matter boundary these borders align preferentially with the sulcal fundi and gyral crowns as all the streamlines seeded from the gyral walls tend to cluster together (Figure 7B). Seeding from the deep/gyral white matter interface eliminates this bias. This reduction of the gyral bias creates a better alignment of the structural connectivity gradients across subjects, which leads to more robust detection of these gradients when averaging across subjects (Figure 9). It also increases the symmetry in tractography with the connectivity estimated by seeding streamlines in a vertex becoming much more similar to the connectivity estimated when considering the streamlines terminating in a vertex (Figure 11).

More promising evidence comes from comparison between the structural and functional connectome for which we show a qualitative improvement in the intrahemispheric connectivity (green arrows in Figure 10) and a small quantitative improvement for the connectivity with the contralateral hemisphere (Figure 11) when adopting the divergence-free model to guide the streamlines through the gyral white matter.

Further validation could come from comparing the connectome estimated from tractography with some known connectivity “ground-truth”, such as that interhemispheric connections are stronger between homotopic than heterotopic regions, which our results suggest. An even stronger validation is a comparison with neuroanatomical tracers in non-human primates. Unfortunately, such ground truth connectivity has been published only at the level of cortical regions, not at the level of individual vertices. Because many of these cortical regions span both sulcal fundi and gyral crowns, the changes in tractography in the gyral blades proposed here has only a minor effect on the parcellated connectomes (Figure S2B,C). Still for completeness, we do include such comparisons in the supplementary materials, where we find that adopting our approach increases the preference for interhemispheric streamlines to connect between homotopic regions (Figure S2D), but find a slightly decreased correlation with tracer data in a macaque dataset (Figure S3).

### 5.3 Alternatives

Explicit constraints on the streamline density like the ones used here to reduce the gyral bias could also be used as part of the cost function in other algorithms. This would not work for local tractography algorithms that only model a single streamline at a time as there is not a meaningful measure of the streamline density. Global tractography algorithms such as the spin glass model (Mangin et al. 2002; Kreher, Mader, and Kiselev 2008; Fillard, Poupon, and Mangin 2009; Reisert et al. 2011) that model all streamlines at once could be used to measure and constrain the streamline density. The spin-glass model might be a better model for U-fibres or other local axons as it allows streamlines within a single bundle to cross each other. Recently, (Teillac et al. 2017) proposed an extension on the spin-glass model to reduce the gyral bias, although their proposal alters the target fibre orientations close to the sulcal walls to allow streamlines to smoothly bend into the gyral walls rather than an explicit constraint on the streamline density. (Wu et al. 2019) also showed a reduction in the gyral bias by encouraging a smooth transition between the radial fibre orientation in the grey matter and the tangential orientation underneath using asymmetric fibre orientation distribution functions (Bastiani et al. 2017).

Density constraints on streamline endpoints could also be added as part of the cost-function, when filtering or weighting streamlines in post-processing (Daducci et al. 2016) by algorithms such as Contrack (Sherbondy et al. 2008), SIFT/SIFT2 (Smith et al. 2013; 2015), LiFE (Pestilli et al. 2014), or COMMIT (Daducci et al. 2015). These algorithms have in common that they filter or assign weights to streamlines produced by local tractography algorithms to represent their relative contribution. While so far these weights are only fitted to the diffusion MRI data, the surface density could be added as an additional constraint. Of course, this does require generating enough streamlines that there is a sufficient population of streamlines connecting to the sulcal walls and fundi. Streamlines connecting sulcal fundi at both ends are so rare (Figure 8) that even after post-processing they might be underrepresented in the final fibre population. Therefore, this post-processing approach might achieve a reduction of gyral bias simply by upweighting the fundi-to-crown connections and not include the many fundi-fundi connections found when tracking to the deep/gyral white matter interface (Figure 8C).

In our approach, the gyral bias is reduced not due to the enforcement of a uniform density across the cortical surface for the vector field, but in using the vector field to map the cortical surface to a less convoluted surface, namely the deep/gyral white matter interface. Tractography to this less convoluted surface does not suffer from a gyral bias. (St-Onge et al. 2018) proposed using a mean-curvature flow model to produce such a less convoluted surface. Their model has the advantage of being much less computationally expensive than the fitting of a vector field to the gyral white matter proposed here. While the reported decrease in the gyral bias seen in St-Onge et al. (2018) is less than found here, this might simply reflect that their final surface is still far more convoluted than the deep/gyral white matter interface adopted here. Ideally, tracer data such as the one shown in Figure 12 would be used to validate the paths proposed by these algorithms through the gyral white matter.

While these alternative algorithms discussed above reduce the gyral bias, the degree of reduction of the gyral bias as shown in Figure 8, has not been shown before. This might increase the accuracy of long-distance connections although perhaps at the cost of losing any information about short-distance connections, in particular those within a gyrus or U-fibres. Code, documentation, and a tutorial for the algorithm proposed in this paper can be found at https://git.fmrib.ox.ac.uk/ndcn0236/gyral_structure and the surface maps displayed are available in the BALSA database (https://balsa.wustl.edu/study/show/0LGM2).

## 6 Acknowledgement

This study was partially funded by the UK EPSRC grant EP/L023067/1/. SJ is supported by the MRC UK (Grant Ref: MR/L009013/1) and a Wellcome Collaborative Award (215573/Z/19/Z), and SS by the Wellcome Trust (217266/Z/19/Z). The Wellcome Centre for Integrative Neuroimaging is supported by core funding from the Wellcome Trust (203139/Z/16/Z). Human data were provided by the Human Connectome Project, WU-Minn Consortium (Principal Investigators: David Van Essen and Kamil Ugurbil; 1U54MH091657) funded by the 16 NIH Institutes and Centers that support the NIH Blueprint for Neuroscience Research; and by the McDonnell Center for Systems Neuroscience at Washington University.

## 7 Supplementary

**Figure S1.**
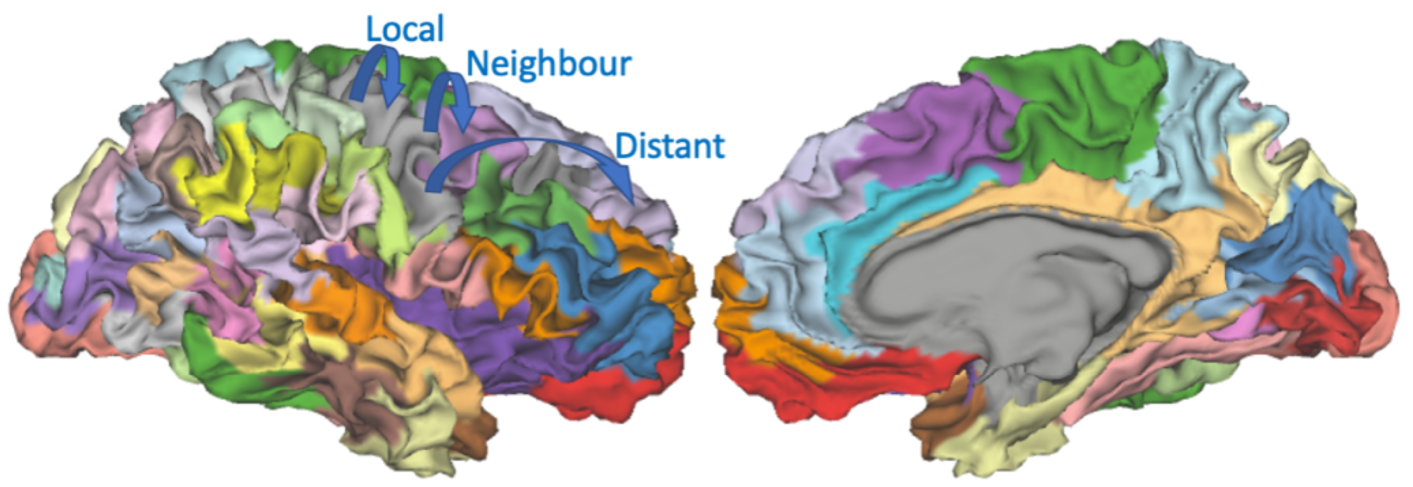
Definition of the intrahemispheric connections excluding local connections and U-fibres used in Figure 11. The cortex is sub-divided into 50 parcels (illustrated for single subject) with the borders between the parcels preferentially located in the sulcal fundi. Local connections and U-fibres are excluded by only considering the connectivity between vertices in different parcels that do not border each other. The parcellation is obtained using a watershed algorithm: the vertices are sorted by sulcal depth (from high to low) and then iterated through. Each vertex is assigned to a new parcel if none of its neighbours are in existing parcels (i.e., it is a local maximum) and otherwise assigned to the neighbouring parcels. If two parcels touch, they are merged if one of them is insufficient deep (i.e., maximum – minimum sulcal depth is below some threshold), otherwise they are kept separate. The depth threshold is chosen, so that we end up with 50 parcels.

**Figure S2.**
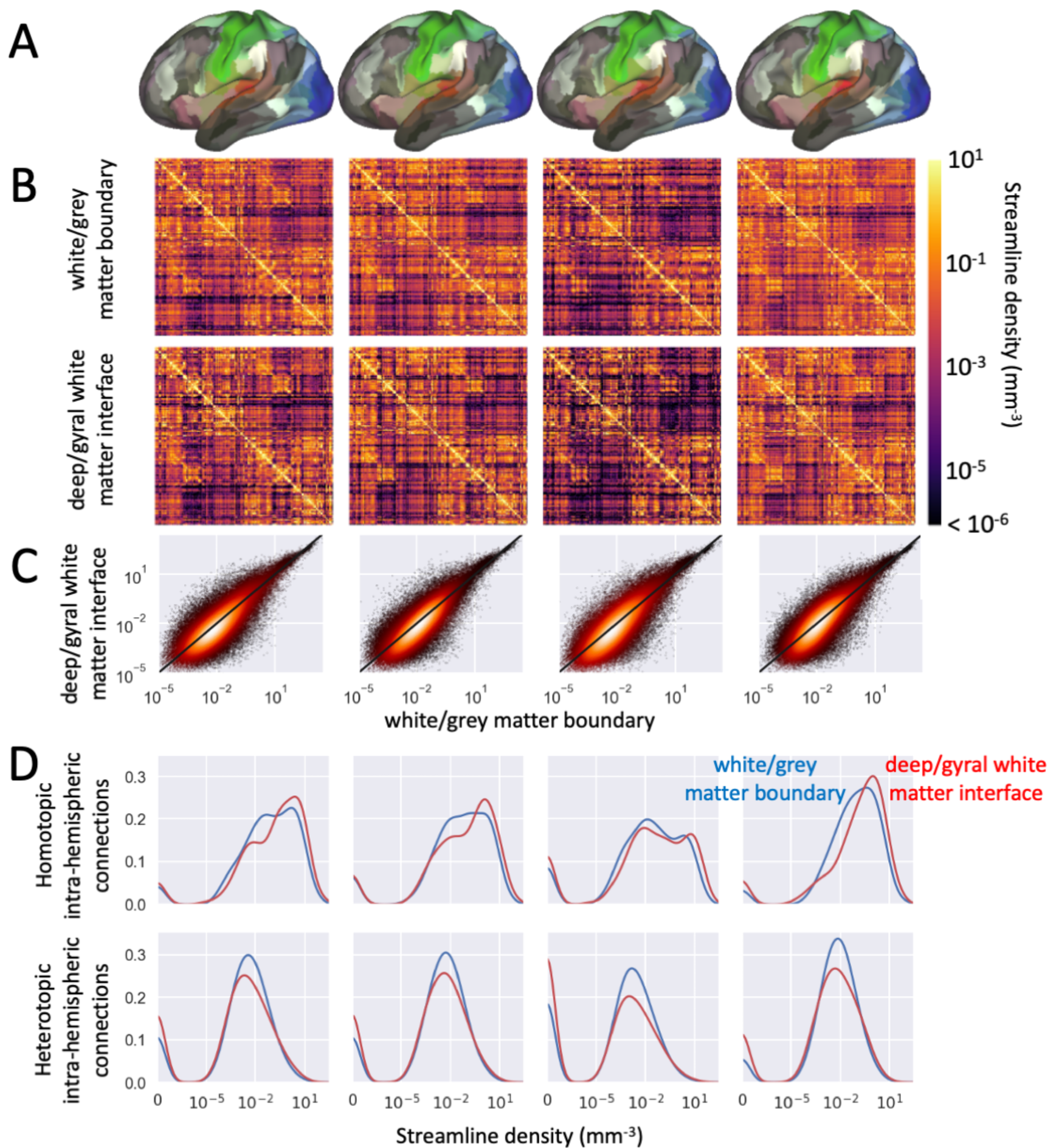
Analysis of parcellated connectomes for 4 subjects (one column per subject). A) Single-subject multi-modal parcellation from Glasser et al. (2016) used to parcellate the connectome. B) Heat map of the streamline density obtained using the white/grey-matter boundary (top) or the deep/gyral white matter interface. The parcellated connnectomes are very similar as also seen in the scatter plot (C). D) Distribution of streamline density for the interhemispheric streamline density between homotopic parcels (top) and heterotopic parcels (bottom). The homotopic connectivity has a median increase of 66%, 37%, 35%, and 53% for these 4 subjects when adopting the deep/gyral white matter interface (red), while the heterotopic connectivity has a median decrease of 10%, 11%, 8%, and 20%. The streamline density measures the number of streamlines seeded from the vertices in one parcel that terminate in each mm^-3^ of the target parcel.

**Figure S3.**
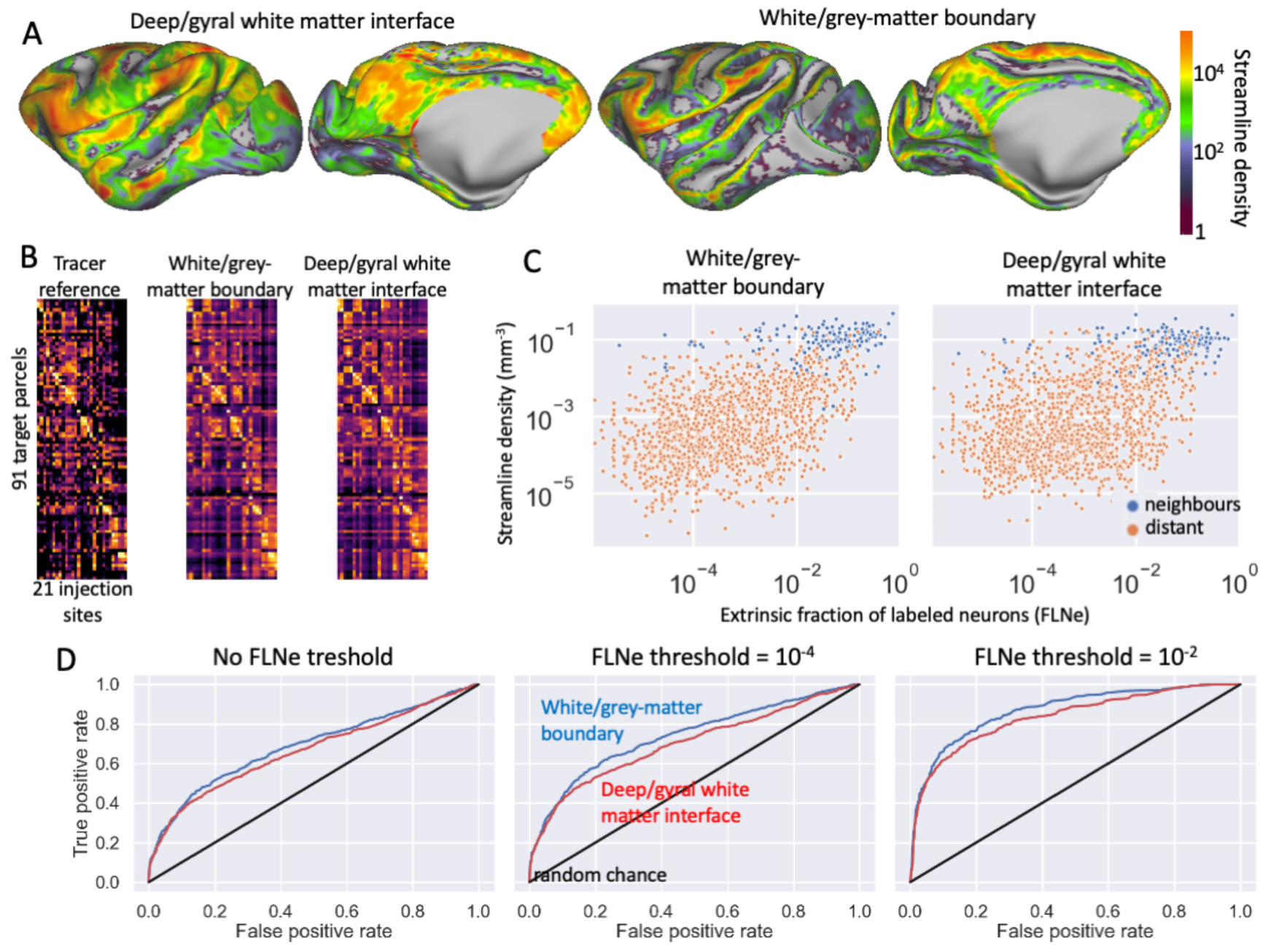
Comparison of the parcellated connectome of a macaque with tracer data from Markov et al. (2011; 2014). A) Illustration of the gyral bias in the macaque diffusion MRI data showing the density of interhemispheric streamline termination points (colour map is the same as in Figure 8A). B) Heatmap of the reference connectome with the log-transformed extrinsic fraction of labelled neurons (FLNe) on the left and the connectomes when using the white/grey-matter boundary in the middle and the deep/gyral white matter interface on the right. C) Correlation between the connectomes from tractography and the tracer connectome (only for the 62% of connections with non-zero connectivity). When adopting the deep/gyral white matter interface the correlation with the tracer connectome becomes worse. This negative trend becomes statistically insignificant when regressing out distance as in Donahue et al. (2016). D) ROC curves for predicting the “true” connections by thresholding the tractography connectomes for from left to right different thresholds of the FLNe. Again, the ROC curves are worse when adopting the deep/gyral white matter interface.

